# Inertial effect of cell state velocity on the quiescence-proliferation fate decision in breast cancer

**DOI:** 10.1101/2023.05.22.541793

**Authors:** Harish Venkatachalapathy, Cole Brzakala, Eric Batchelor, Samira M. Azarin, Casim A. Sarkar

## Abstract

Energy landscapes can provide intuitive depictions of population heterogeneity and dynamics. However, it is unclear whether individual cell behavior, hypothesized to be determined by initial position and noise, is faithfully recapitulated. Using the p21-/Cdk2-dependent quiescence-proliferation decision in breast cancer dormancy as a testbed, we examined single-cell dynamics on the landscape when perturbed by hypoxia, a dormancy-inducing stress. Combining trajectory-based energy landscape generation with single-cell time-lapse microscopy, we found that initial position on a p21/Cdk2 landscape did not fully explain the observed cell-fate heterogeneity under hypoxia. Instead, cells with higher cell state velocities prior to hypoxia, influenced by epigenetic parameters, tended to remain proliferative under hypoxia. Thus, the fate decision on this landscape is significantly influenced by “inertia”, a velocity-dependent ability to resist directional changes despite reshaping of the underlying landscape, superseding positional effects. Such inertial effects may markedly influence cell-fate trajectories in tumors and other dynamically changing microenvironments.

## Introduction

Cell fate determination is a complex process, underpinned by nonlinear interactions among various biomolecular components that can obscure intuition of key modulators of cell behavior. Energy landscapes can provide graphically intuitive representations of cell decision-making^1,2^. In this framework, cells are considered as balls rolling on a landscape where each position represents a cell state and the depth at that position represents its potential energy or relative stability. Cell “movement” on the landscape is determined by its initial state, the surrounding topography, and the noisiness of the biomolecular components and interactions within the cell. This framework can also offer quantitative or semi-quantitative descriptions of important landscape features such as valley positions, relative state stability, and mean transition paths and times between these states. Landscape formalisms have been previously used to understand cell decision-making in contexts such as stem cell differentiation^2–5^, embryonic development^6,7^, cell cycle progression^8,9^, DNA damage responses^10,11^, and cancer^12–15^. These studies recapitulate the overall population heterogeneity and dynamics as validated through ensemble-level measurements such as flow cytometry, immunostaining, or single-cell RNA quantification. However, the extent to which individual cell dynamics *in vitro* or *in vivo* are faithfully recapitulated on these landscapes is unclear, particularly under external perturbations such as growth factors, cytokines, or stress. Important features such as the contributions of initial position of individual cells and noisiness of movement on this landscape are often obscured due to a lack of single-cell tracking. For example, it is typically assumed that heterogeneity in phenotype transitions under an external cue is primarily driven by short-term stochastic fluctuations due to intrinsic biochemical noise. However, this is highly context dependent as studies show more persistent, but still stochastic, differences in cell state such as epigenetic variation can also lead to divergent responses^16,17^. Dissecting the specific contributions of these factors is key to engineering cell behavior in different contexts.

In this study we aim to understand how both position and velocity regulate cellular movement on an energy landscape under an external perturbation. Here, we focus on the proliferation-quiescence decision in breast cancer dormancy. We chose this clinically important cell decision because it is governed by a well-established but nonlinear protein network implicated in the dynamic cell cycle, and therefore we hypothesized important roles of cell position and velocity on the landscape for this system. Furthermore, to our knowledge, the effects of stochasticity – changes in cell state or response due to intrinsic fluctuations in the biochemical processes driving cellular behavior – remain unexplored in this fate decision.

Clinically, cancer dormancy presents a significant challenge in progression-free survival in breast cancer patients even after several years of being in complete remission^18^. This is underpinned by the reactivation of dormant breast cancer cells that persist post-treatment, often in hypoxic niches such as the bone marrow^19–21^. These dormant cells arise when typically proliferative tumor cells are faced with stressful conditions, such as hypoxia or nutrient deprivation, causing them to enter a dormant state characterized by quiescence, a reversible non-dividing “resting phase”^22^. Notably, tumor cells exhibit significant heterogeneity in their abilities to enter a dormant state^22–24^. This variability is thought to be driven by genetic differences (e.g., estrogen receptor (ER) expression^25^) and/or cell states (e.g., stem-cell-like phenotypes^26,27^). However, there is heterogeneity in dormancy even among cells with a given dormancy-associated phenotype^23,27,28^. It is unclear whether this is purely due to microenvironmental differences or if there are also stochastic determinants arising from intrinsic biomolecular fluctuations. Strikingly, heterogeneity in quiescence induction, a core feature of dormancy, has been shown in some contexts to be largely recapitulated by a toggle switch consisting of the cyclin-dependent kinase (CDK) Cdk2 and its mutually antagonistic partner p21^29–33^, a CDK inhibitor shown to be upregulated in some dormant tumor phenotypes^20^. We thus hypothesized that analyzing variability in quiescence induction under hypoxia through this core p21-Cdk2 motif – specifically, the contributions of position (biochemical activities/expression of p21 and Cdk2) and velocity (change in cell state defined by these species) to cell behavior – can provide insight into the non-genetic heterogeneity in dormancy induction.

Applying a trajectory-based energy landscape methodology^34^, supported by live-cell fluorescence microscopy and single-cell tracking, we mapped the underlying p21-Cdk2 landscape under normoxic and hypoxic conditions in MCF-7 breast cancer cells, finding highly variable quiescence-proliferation cell behavior within the population. We examined the dependence of cell fate on initial position and velocity, surprisingly discovering heterogenous cell behavior to be primarily driven by differences in cell state velocity while initial position was only partially explanatory of decision-making in this context. We observed that the cells exhibit “inertia” – a tendency to continue moving along a direction of motion unless acted on by an external force. In this context, proliferative cells preferred to remain proliferative unless the change in landscape due to hypoxia was strong enough to change their direction of movement and push them towards a quiescent state. Finally, we demonstrated the epigenetic inheritability of this phenomenon, showcasing additional system information encoded in velocity, compared to position alone.

## Results and Discussion

### Single-cell heterogeneity in cell cycle response to hypoxia in MCF-7 cells can be explained by a p21-Cdk2 potential energy landscape

Tumor cells have differential abilities to enter a dormant state depending on genotypic and phenotypic features.^22,25–27^ For example, ER^+^ cells typically exhibit longer periods of dormancy before relapse than ER^-^ cells do.^25^ However, individual ER^+^ tumor cells can also differ in their ability to enter a dormant state.^25,27,28^ While this can be due to microenvironmental factors, stochastic fluctuations in the pathways governing dormancy may also contribute. To examine this contribution, we used a hypoxia-mimetic CoCl_2_ platform that has previously been shown to recapitulate the heterogeneity in dormancy capabilities associated with the ER^+^ MCF-7 and ER^-^ MDA-MB-231 breast cancer cell lines *in vitro* in addition to single-cell heterogeneity within MCF-7 cells^28^. This platform provides a uniform microenvironment, minimizing extrinsic variability, and can be used to draw conclusions about cell-to-cell heterogeneity due to genetic and epigenetic variability as well as intrinsic noise. Further, the introduction of CoCl_2_, hereafter referred to as hypoxia, is convenient and more stable compared to conventional hypoxic chambers wherein the normoxic response is established rapidly whenever cells are removed for manipulation and analysis. Additionally, in contrast to true hypoxia where GFP variants exhibit reduced fluorescence due to improper folding, CoCl_2_-based hypoxia can be used with a wider range of non-overlapping fluorophores^35,36^ allowing for simultaneous marker tracking in live-cell imaging experiments.

Identifying the drivers of heterogeneous dormancy behavior under hypoxia in MCF-7 cells is challenging due to the number of phenotypic features associated with dormancy. However, quiescence, a reversible exit from the cell cycle into the G_0_ phase, is a conserved feature of cellular dormancy implying that heterogeneity in the levels of p21 and Cdk2, drivers of quiescence and proliferation, respectively, can be used to understand heterogeneity in dormancy capacities. This is consistent with previous observations of a mutually antagonistic p21-Cdk2 toggle switch underpinning quiescence induction in single cells.^30–32,37,38^ Specifically, high Cdk2/low p21 states are proliferative while low Cdk2/high p21 states are quiescent (**Figure 1A**). Indeed, by western blotting, we found that MCF-7 cells showed upregulation of p21 along with downregulation of Cdk2 under hypoxia (**Figure 1B**). Further, we did not see any changes in p21 or Cdk2 expression in MDA-MB-231 breast cancer cells that do not enter dormancy under hypoxia (**Supplementary Figure 1**), validating the use of this motif to examine dormancy regulation.

**Figure 1:**
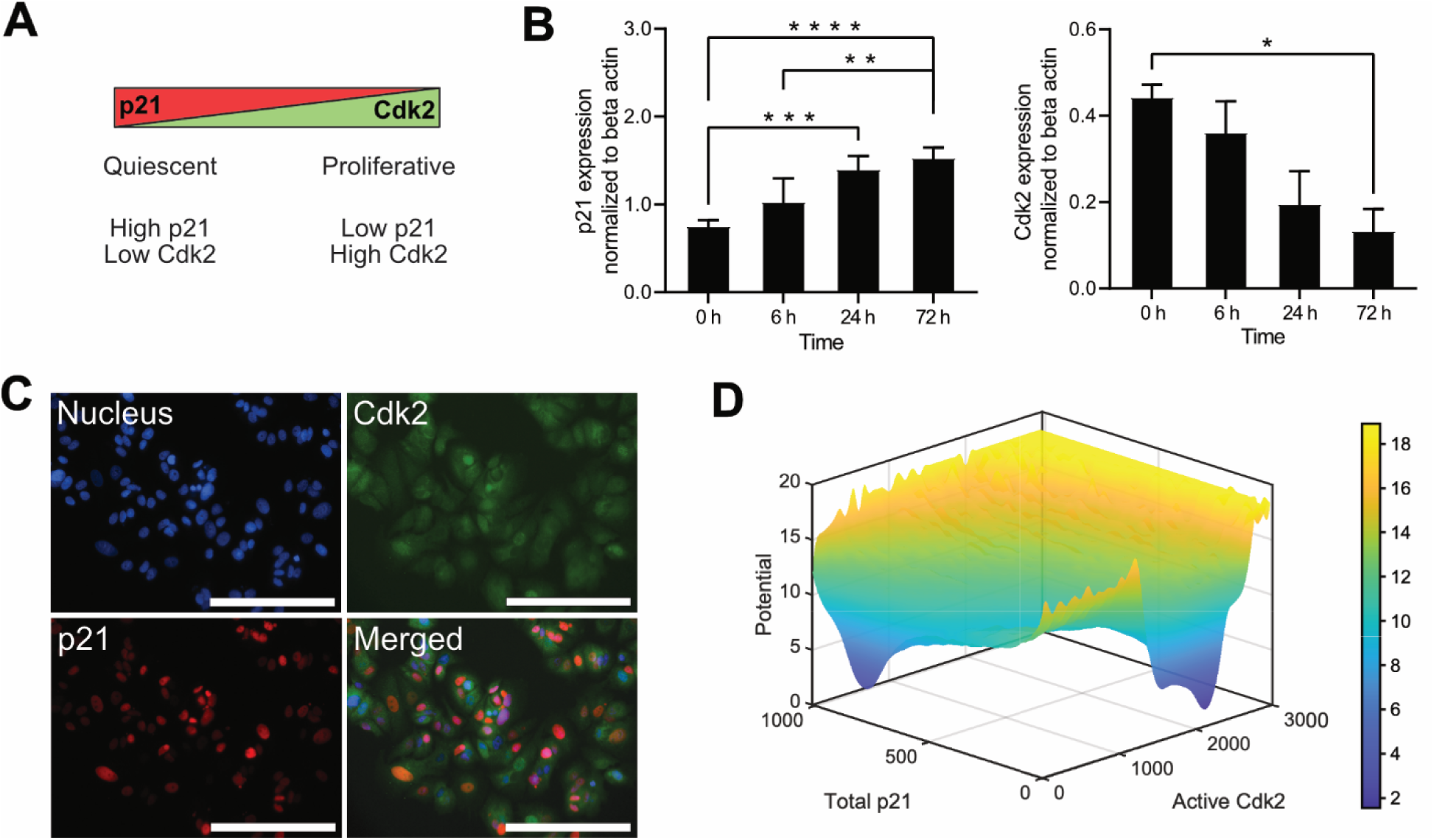
Heterogeneity in quiescence-proliferation under hypoxia in MCF-7 cells is explained by a p21-Cdk2 toggle switch. **(A)** Schematic of the p21-Cdk2 dependence of the quiescence-proliferation decision **(B)** Quantification of western blots of p21 (left) and Cdk2 (right) expression under cobalt chloride treatment in MCF-7 cells at different times. Bar graphs and error bars represent the mean and s.d. of three independent experiments. p-values calculated by ANOVA followed by Dunn’s post-hoc test with multiple comparison corrections (* p < 0.05, ** p < 0.01, *** p < 0.001, **** p < 0.0001) **(C)** Immunostaining images of MCF-7 cells under hypoxia showing nuclei stained by DAPI (top left), Cdk2 (green, top right), p21 (red, bottom left), and merged (bottom right). Scale bar represents 200 µm. Image brightness and contrast adjusted for clarity. **(D)** Potential energy landscape of the p21-Cdk2 computational model as a function of total p21 and active Cdk2.

We then examined whether the p21 and Cdk2 motif remains explanatory of the quiescence-proliferation decision in this context at a single-cell level. Using immunofluorescence microscopy, we observed significant heterogeneity in p21 and nuclear Cdk2 expression under hypoxia with minimal evident co-expression of p21 and Cdk2, supporting the notion of a mutually exclusive toggle switch between the two proteins (**Figure 1C**). We inferred two subpopulations which are defined as p21^high^/Cdk2^low^ (quiescent) and p21^low^/Cdk2^high^ (proliferative). Further, we observed a strong positive correlation between nuclear Cdk2 and the Ki67 proliferation marker intensities under both normoxic and hypoxic conditions **(Supplementary Figure 2).** Taken together, these results validate the presence of a p21-Cdk2 toggle switch motif that underlies the single-cell quiescence-proliferation decision in MCF-7 cells under hypoxia.

We then sought to examine whether such phenotypic heterogeneity can be explained by the underlying energy landscape. To this end, we applied a trajectory-based landscape generation methodology^34^ to the stochastic version of a quiescence-proliferation model developed by Heldt et al.^38^; however, the original model describes the S phase as a transitory state and the G_2_ phase as the final steady state, distracting from the quiescence-proliferation decision itself, which is the decision of interest on the landscape. Thus, for this analysis, the model was modified to exclude the exit from S phase to focus solely on the decision to enter either S or G_0_. The underlying landscape (**Figure 1D**) generated using trajectories with randomly initialized abundances of total p21 and active Cdk2, shows the existence of two valleys – p21^high^/Cdk2^low^ and p21^low^/Cdk2^high^. These represent the stable steady states of the system as shown by Heldt et al. and correspond to the subpopulations observed by immunostaining where p21^high^/Cdk2^low^ indicates quiescence and p21^low^/Cdk2^high^ indicates proliferation. In terms of cell cycle progression, the p21^high^/Cdk2^low^ state refers to G_0_, the p21^low^/Cdk2^high^ state refers to S phase, and the transitory region between these states is the G_1_ phase.

A similar landscape with the same mathematical model, constructed by a different methodology, was used by Chen and Li^9^ to quantify the mean multi-dimensional paths involved in quiescence-proliferation transitions. In contrast with our study, which uses parameters identified in Heldt et al., their landscape used modified parameters such that three stable steady states exist – G_0_ quiescence, G_1_ cell cycle arrest, and S phase. While the landscapes are qualitatively similar in appearance, the striking difference in our analyses is the characterization of the G_1_ phase as either a steady state on the Chen and Li landscape or a transitory state on our landscape. However, previous single-cell studies on the cell cycle indicate cells do not spend large amounts of time arrested in G_1_ and instead enter G_0_ when faced with stress, supporting the transient nature of the G_1_ phase; this is evidenced by the lack of a distinct valley in between the quiescent p21^high^/Cdk2^low^ and proliferative p21^low^/Cdk2^high^ valleys on our landscape^30,31,39,40^. This discrepancy arises due to the methodologies used in each case. Chen and Li use a mean-field approximation-based approach to calculate the steady state probability distributions of the system at equilibrium, necessitating the G_1_ state to be a steady state to appear as a valley on the landscape. In contrast, the trajectory-based method^34^ can capture intermediate transition states such as the G_1_ phase without the need for recharacterization as a steady state. Overall, our results imply that heterogeneity in the quiescence-proliferation decision can be explained by an underlying bistable energy landscape computed using the trajectory-based method.

### Hypoxia biases the energy landscape towards a quiescent phenotype, but still permits heterogeneity that is not fully explained by position alone

Having validated the presence of a p21-Cdk2 energy landscape explanatory of cell behavior in this context, we sought to qualitatively map the change in landscape topography when switching from normoxic to hypoxic conditions. From the population-level data of p21 and Cdk2 expression in MCF-7 cells under hypoxia (**Figure 1B**), we inferred that cells upregulate p21 and simultaneously downregulate Cdk2. p21 upregulation in this context is likely driven by the stabilization of its transcription factor p53 under hypoxia^41^. This was recapitulated in the model by decreasing the p53 degradation rate. Similarly, simultaneous Cdk2 downregulation can be driven by p53-mediated *MYC* transrepression, an interaction not modeled in the original computational framework but found to be present in MCF-7 cells^42^. We modeled this interaction as a reduction in mitogenic signaling by reducing the basal synthesis rate of the transcription factor E2f.

The transition from normoxia to hypoxia caused the cell-fate bias on the energy landscape to transform from strongly proliferative to strongly quiescent, as evidenced by the change from a deep proliferative valley at high Cdk2/low p21 and shallow quiescent valley at low Cdk2/high p21 to a shallow proliferative valley and deep quiescent valley (**Figure 2A**). Interestingly, we also observed a small valley at a low Cdk2/low p21 state under hypoxic conditions representing a metastable state wherein cells were driven to quiescence primarily by a lack of mitogenic signaling as opposed to increased p21 upregulation (**Figure 2A**). Our simulations suggested that this state can be a stable steady state that arises when p21 synthesis is low and mitogenic signaling is not sufficiently high for the cells to proceed further in the cell cycle (**Supplementary Figure 3**); this is akin to a serum starvation-induced p21^low^/Cdk2^low^ quiescent phenotype previously reported by Barr et al.^31^ However, due to the expected upregulation of p21 under hypoxia, we predict this state to be a transitory state on the path to quiescence as opposed to being a more persistent quiescent state.

**Figure 2:**
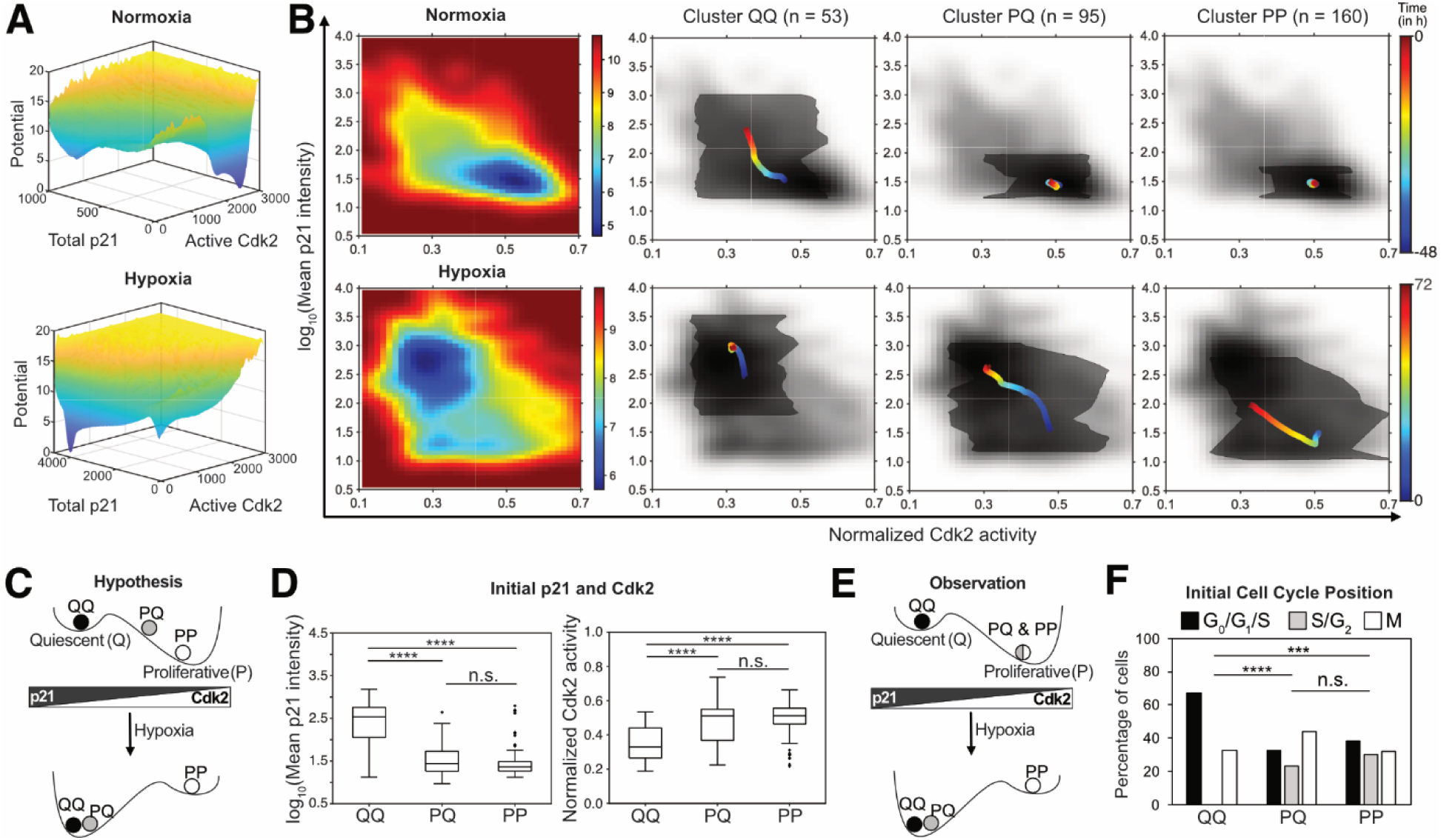
Energy landscapes under normoxia and hypoxia and the role of position on the observed cell-fate divergence. **(A)** Computationally calculated energy landscapes under normoxia (E2f synthesis rate = 1.19 (i.e., 2^0.25^) times the basal value) and hypoxia (E2f synthesis rate = 0.71 (i.e., 2^-0.5^) times and p53 degradation rate = 10^-2^ times the basal values). **(B)** Experimental energy landscapes (left) generated using single-cell tracking imaging data and the subpopulations of behavior (right) observed under normoxia followed by hypoxia: cells entering a quiescent state in normoxic conditions that remain quiescent under hypoxia (Cluster QQ), cells proliferative under normoxia but quiescent under hypoxia (Cluster PQ), and cells proliferative under normoxia that remain proliferative under hypoxia (Cluster PP). **(C)** Pictographic representation of the hypothesized dependence of cell fate on initial position. **(D)** Effect of initial p21 levels and Cdk2 activities on cell fate under hypoxia. Statistical comparison of mean ranks was carried out by Kruskal-Wallis test followed by Bonferroni correction for multiple comparisons. **(E)** Pictographic representation of the observed dependence of cell fate on initial position. **(F)** Effect of initial cell cycle position on cell fate. Distributions were compared using the Kolmogorov-Smirnov test for discrete distributions followed by Bonferroni correction for multiple comparisons. (** p < 0.01; *** p < 0.001; **** p < 0.0001).

To validate our computational predictions, we used time-lapse imaging on MCF-7 cells engineered to express p21-mVenus from the native *CDKN1A* (gene encoding p21) genomic locus, DHB-mCherry (a Cdk2 sensor)^30^, and a H2B-mTurquoise nuclear marker and subjected them to hypoxia through CoCl_2_ treatment. Using single-cell tracking and fluorescence quantification, we obtained p21 expression and Cdk2 activity levels for individual cells, imaged every 20 minutes, over a period of 120 hours (48 hours under normoxia, 72 hours under CoCl_2_). We used these experimental p21 and Cdk2 trajectories to map the underlying steady-state landscape under normoxia and hypoxia. In agreement with our simulations, we observed a strong reduction in the permissivity for proliferation in the landscape under hypoxia characterized by a shallow proliferative valley and deep quiescent valley (**Figure 2B**). We also identified a small Cdk2^low^/p21^low^ valley under hypoxia, but it is not clear from experimental data alone whether this is a transitory state or a more permanent quiescent state. Longer term experiments (∼1-2 weeks) would be required to improve confidence in the characterization of this state. Interestingly, coexistence of the p21^low^/Cdk2^low^ and p21^high^/Cdk2^low^ quiescent phenotypes highlights the redundancy in quiescence induction driven by simultaneous inhibition of mitogenic signaling (lower E2f synthesis rate resulting in lower Cdk2 activation) and upregulation of stress signaling (lower p53 degradation rate resulting in higher p21 expression). This redundancy can aid in more robust quiescence induction as well as in the maintenance of this state, particularly if one of the pathways is more prone to noise. Further, this can result in increased heterogeneity in the biomolecular identities of the quiescent states and improved survival when specific pathways are targeted. There exist analogous cases in proliferative cancer cells where either the redundancies in pathways^43,44^ or the heterogeneity in biomolecular states^45,46^ can result in maintenance or recovery of a proliferative phenotype.

In general, the cells transited from the proliferative valley to the quiescent valley under hypoxia as evidenced by the shift in landscape permissivity. However, given the presence of multiple valleys in the hypoxic landscape, we hypothesized that cells would exhibit divergent p21-Cdk2 dynamics when the landscape is perturbed due to hypoxia. Using unsupervised clustering (see **Materials and Methods**), we identified four clusters of p21-Cdk2 dynamics (**Supplementary Figure 4**). Comparing the clusters, we observed qualitatively different types of behavior for each with respect to the quiescence-proliferation decision immediately after hypoxia: (a) Cluster QQ – cells that either entered or started to enter quiescence before hypoxia treatment, remaining quiescent under hypoxia, (b) Cluster PQ – cells that entered quiescence under hypoxia treatment, (c) Cluster PP – cells that continued along the cell cycle under hypoxia treatment (**Figure 2B**). The fourth cluster of cells showed similar behavior to cluster QQ with cells exhibiting high p21 and low Cdk2 levels under both normoxic and hypoxic conditions; classifying the data into three clusters, instead of four, showed that this fourth cluster merged with cluster QQ (**Supplementary Figure 4C, D**). However, this cluster had significantly fewer cells (n = 6) compared to the other clusters (n = 53, 95, and 160) and, therefore, it was discarded in order to provide better statistical significance for inter-cluster comparisons moving forward. We also note that cells in cluster PP eventually entered quiescence at variable points of time after the first fate decision, either with or without cytokinesis. However, the analysis of such longer-term behavior is outside the scope of this work.

Since dynamics and cell fate on the computational landscape are influenced by the initial cell position, we hypothesized that variability in initial position defined by p21 abundance and Cdk2 activity could be a potential source of the observed cell fate heterogeneity. We investigated whether initial p21 expression and Cdk2 activity are predictive of the inter-cluster differences under hypoxia. p21 expression would be expected to be higher in QQ and PQ in comparison to PP, while Cdk2 activity would be expected to be lower in QQ and PQ in comparison to PP (**Figure 2C**). At the onset of hypoxia (t = 0 h), cluster QQ showed higher initial p21 expression and lower initial Cdk2 activity compared to clusters PP and PQ (p < 0.0001; Wilcoxon Rank Sum Test); however, there were no statistically significant differences in initial p21 expression or initial Cdk2 activity between clusters PP and PQ, despite the divergence in cell fate (**Figure 2D, E**).

The degeneracy of the clusters with respect to the initial p21 expression and Cdk2 activity could be reflective of the complexity of the quiescence-proliferation decision. While p21, Cdk2, and the two-dimensional energy landscape involving these species are ultimately reflective of the cell fate in this context, there are several other biochemical species such as Rb and Cyclin D involved in the cell decision-making process whose instantaneous levels are not represented by p21 and Cdk2 alone. The instantaneous levels of all these species determine the initial position of the cells on the multi-dimensional landscape describing cell fate and can lead to divergent cell fate responses even with similar p21 and Cdk2 levels. However, it is impractical to simultaneously obtain temporal information *in situ* on all the species whose instantaneous levels could affect downstream decision-making. Instead, we relied on the dependence of these various species on cell cycle position and used Cdk2 activity dynamics as a qualitative proxy for cell position on the high-dimensional landscape. Using the relative shape of Cdk2 activity traces aligned to cell division, we classified the cells into either G_0_/G_1_/S, S/G_2_, or M (**Materials and Methods**). While there was a difference in the cell cycle position distribution between cluster QQ and PQ as well as between QQ and PP (p < 0.0001 and p < 0.001 respectively; Kolmogorov-Smirnov test), PP and PQ did not have statistically different distributions (**Figure 2F**). While our analysis did not use more precise cell cycle tracking markers such as the FUCCI sensors^47^, we can still conclude that cells that remained proliferative under hypoxia were not solely cells that were in a different phase of the cell cycle.

For cells that divided after hypoxic treatment, one daughter cell was randomly picked for cell fate analysis to avoid multiple counting of the same cell. Including all daughter cells in the analysis did not change the results significantly; the conclusions drawn from the data remain the same (**Supplementary Figure 5A, B**). Additionally, controlling for approximate cell cycle phase when considering initial position did not change the conclusions drawn from the results (**Supplementary Figure 6**). Overall, our analysis demonstrates that initial position on the landscape, determined either by p21-Cdk2 levels or by the approximate cell cycle position, is unable to fully explain the observed inter-cluster quiescence-proliferation heterogeneity.

### Cell state velocity displays an inertial effect on the quiescence-proliferation decision

Previous studies show that flux is a determinant of cell fate in addition to position in non-equilibrium systems^8,10,48^. Given the dynamic nature of the cell cycle, we hypothesized that cell state velocity – that is, the rate of change of the biochemical determinants of the cell state – may be more determinative of cell fate, especially when the landscape is perturbed. This velocity, while a function of position, is also determined by the parameters governing the system, such as rate constants in an ODE-based description. These parameters represent critical effectors of cell fate, such as epigenetic regulation and signaling crosstalk, that are not explicitly described by the abundances of the system variables (i.e., the position on the landscape). Thus, velocity as a cell decision-making criterion can provide key information related to the system that is not encompassed by a multi-dimensional position that includes all model variables.

Since the cell cycle is a dynamic system, we used cell cycle velocity as a measure of the average cell state velocity. In the mathematical model, cell cycle velocity was defined as the inverse of time to S phase entry. Experimentally, we used the number of cell divisions in the imaging period 48 hours prior to hypoxic treatment as an analogous cell cycle velocity measure. First, using a local sensitivity analysis on the mathematical model, we determined the top ten system parameters that affect time to S phase entry. We found these parameters also ranked within the top ten in overall local sensitivity (as performed by Heldt et al.^38^), suggesting that parameters affecting cell cycle velocity could also be important for the quiescence-proliferation decision (**Figure 3A**). Additionally, parameter changes resulting in slower cell cycle velocities also resulted in a shift from proliferation to quiescence (**Supplementary Figure 7**), suggesting that cells with slower cell cycle velocities would be more likely to enter quiescence. Indeed, we found a positive correlation between the number of cell divisions prior to hypoxia and the percentage of cells in cluster PP, suggesting an increased propensity to remain proliferative under hypoxic stress in faster cells (**Figure 3B**). In other words, cell cycle velocity has an inertial effect on cell fate, wherein faster cells have a higher propensity to continue along their path despite an external “force,” arising from topographical changes in the landscape due to environmental perturbations, that biases against the direction of motion. In contrast, slower cells are more likely to change their direction of motion in response to this force and enter a different state.

**Figure 3:**
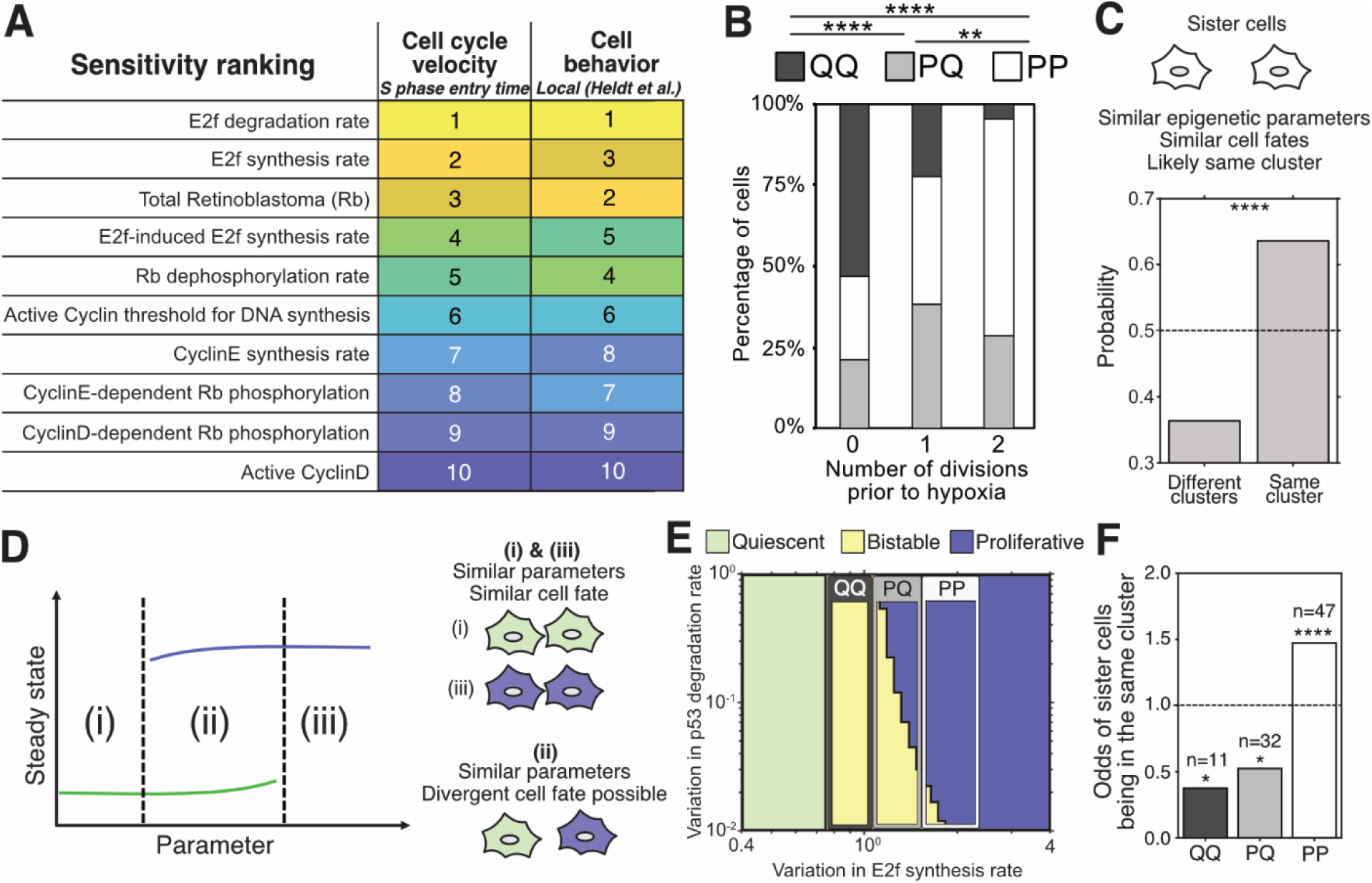
System parameters drive the inertial effect of velocity. **(A)** Ranking of system parameters based on S phase entry time sensitivity analysis in this work and local sensitivity analysis by Heldt et al. showing the commensurate effect of parameter variations on cell cycle velocity and cell behavior, respectively. **(B)** Distribution of cell cluster identity as a function of number of cell divisions prior to hypoxia. Division number distributions between different clusters were compared with a discrete version of the Kolmogorov-Smirnov test. **(C)** Probabilities of sister cells being in the same cluster or different clusters. Statistical significance is calculated by 10^5^ instances of probability calculation after randomization of cluster identities. **(D)** Schematic showing the effect of system bistability on cell fate. Cells in either extreme of the parameter space (i and iii) are likely to have the same fate, whereas cells either in the bistable region (ii) or at the cusp of bistability can exhibit divergent cell fates due to stochastic effects. **(E)** Phase space diagram showing the dependence of cell fate on the deviation of p53 degradation rate and E2f synthesis rate from basal values. Green and violet regions represent quiescence and proliferation, respectively. Yellow region represents a bistable region. The expected parameter ranges for each cluster are shown as hollow rectangles. Graph recolored and boundaries drawn post-analysis for clarity **(F)** Odds of the sister cell of a cell in clusters QQ, PQ, or PP being in the same cluster (for a given cell, probability of its sister cell belonging to the same cluster/probability of its sister cell belonging to a different cluster). Statistical significance was calculated by 10^5^ instances of probability calculation after randomization of cluster identities. (* p < 0.05; ** p < 0.01; *** p < 0.001; **** p < 0.0001).

Nathans et al.^40^ reported a similar inertial effect reflective of the delay in p21 upregulation post-DNA damage. In their study, “older” cells that were further along G_1_ were more likely to enter S phase despite DNA damage as they had less time to upregulate p21 sufficiently to stop entry into the S phase. In contrast, “younger” cells that were earlier in G_1_ were more likely to enter quiescence. However, we observed no such distinction in this study; cells that remained proliferative were not necessarily “younger”. There was no statistically significant difference in the age distributions of quiescent and proliferative cells relative to hypoxia induction (**Supplementary Figure 8**). Previous studies also showed that stochastic DNA damage in the mother cell cycle can lead to longer cell cycles and increased propensity for daughter cell quiescence, driven by p21 upregulation starting in the M phase of the mother cell^31,49^. While we observed higher p21 expression in cells that are in the M phase at the time of hypoxic treatment in cluster QQ, there were no statistically significant differences between clusters PQ and PP (**Supplementary Figure 6**). Taken together, these findings suggest that the inertial effect of cell cycle velocity observed in our study is likely driven by other biochemical or biophysical properties. Given that velocity is influenced by system parameters, some of which are determined by epigenetics, we hypothesized that the effect of velocity on cell fate exhibits inheritability. In other words, we expect that the subsequent cell fate decision should be similar between daughter cells. By examining the cluster identity of sister cells born before hypoxia, we found that there is a high probability (P = 0.64; p < 0.0001) of sister cells belonging to the same cluster (**Figure 3C**). In contrast, the mean probability of sister cells being in the same cluster for a randomized dataset with the same distribution of cluster identities was much lower at 0.36 (**Supplementary Figure 9A**). Interestingly, 36% of sister pairs had divergent cell fates (**Figure 3C**). This can be explained by the presence of stochasticity in system parameters as well as bistability within the system. Sister cells with parameters within the bistable region or close to it can have divergent cell fates when combined with small changes in system parameters or stochastic dynamics (**Figure 3D**). We examined the phase space of the p21-Cdk2 dynamical system using basal E2f synthesis and p53 stability as proxies for mitogenic and stress signaling, respectively. With increasing E2f synthesis rates (corresponding to enhanced mitogenic signaling), we observed a transition from monostable quiescence to monostable proliferation bridged by a bistable region of varying widths; the width of this bistable region is dependent on the p53 degradation rate (with lower degradation rates corresponding to increased stress signaling), suggesting the potential for such heterogeneity to arise. Through this phase space analysis and the general proclivity of cancer cells towards proliferation, we expected proliferative cells (cluster PP) to be in the monostable proliferative region while cells becoming quiescent in our study (clusters QQ and PQ) were expected to be within the bistable region, where cells can be both quiescent and proliferative (**Figure 3E**). This implied clusters QQ and PQ were likely to contain cells whose fate is divergent from their siblings while those in cluster PP would have a similar fate as their sibling. Indeed, in our dataset, the odds of both sister cells being in the same cluster for clusters QQ and PQ was less than 1 (p < 0.05) while cluster PP showed a ratio greater than 1 (p < 0.001) (**Figure 3F, Supplementary Figure 9B**).

Analogously, we expect cells with slower velocities to have more heterogeneity in cell fate. To examine this, we use Shannon entropy *E* = -Σ *p_i_·ln(p_i_*), where *p_i_* is the probability of event *i*. A higher entropy implies multiple events are similarly likely to occur, resulting in increased heterogeneity. In contrast, a lower entropy represents a narrower distribution where one event is more likely than others. Using this metric, we found that cells with 0 and 1 divisions prior to hypoxia had high distribution entropies (0.46 and 0.47, respectively) that are approximately equal to the entropy of a uniform distribution with the three bins (0.48); by contrast, cells with 2 divisions had a lower entropy (0.32) (**Supplementary Figure 10**). Additionally, since changes in cell cycle velocity through system parameters are likely to change the dynamics and steady state of the system, we hypothesized that the mean p21 expression is negatively correlated to cell cycle velocity while mean Cdk2 activity is positively correlated to this velocity. By simulating multiple instances of the mathematical model, we found that this correlation exists as hypothesized (**Supplementary Figure 11A, D**) and was qualitatively recapitulated in the experimental data (**Supplementary Figure 11B, E**). Taken together, these results point towards the epigenetic nature of the effects of velocity in this study.

In the context of cancer dormancy, these results suggest that the velocity of the system can influence the quiescence-proliferation decision in individual cells. However, over longer times, we observe that most cells in cluster PP eventually become quiescent after prolonged G_2_ arrest, with or without cell division (**Supplementary Figure 12**), implying that chronic hypoxia can likely overcome the effects of velocity. It is likely that the effects of velocity are more pronounced under transient or fluctuating hypoxia wherein subpopulations of cells can enter quiescence depending on their velocity while others remain proliferative. This hypothesis is challenging to validate using our current experimental system due to the retention of cobalt chloride in cells after washout as well as the misfolding of GFP variants used in live-cell imaging under true hypoxia. However, similar time-dependent phenomena are seen in other systems with toggle switch motifs such as quiescence exit^50,51^, cell differentiation^52^, and synthetic genetic circuits^53^ where heterogeneity in initial biochemical states within the governing bistable switch is amplified leading to divergent phenotypes. In these studies, cells undergo phenotypic transitions at different times leading to variability in cell states under transient or weak cues, similar to cell behavior under early hypoxia observed in our system. In comparison, cells exit quiescence more uniformly or differentiate better under sustained or stronger cues in these examples, analogous to behavior in chronic hypoxia. Together, these lend support for the role of velocity in transient or fluctuating hypoxia in propagating single-cell heterogeneity in dormancy.

Interestingly, our results suggest that velocity and position together serve as a better predictor of cell fate than position alone. Given that velocity is typically determined by the position on an energy landscape, this could point to the inadequacy of p21 and Cdk2 alone in sufficiently determining the position on the overall higher dimensional landscape, implying measuring position using additional biochemical species might be required to better describe cell behavior. For example, Yang et al.^29^ have shown that levels of p53, p21, phospho-ERK, and CyclinD mRNA in a mother cell under stress, albeit not measured simultaneously in a single cell, were predictive of quiescence entry in the daughter cell and are likely to simultaneously influence the fate decision. However, it can be increasingly impractical to simultaneously measure all species that contribute to a cell fate decision. Furthermore, these measurements lack information about the rate constants and other parameters in the system. While this is less concerning for relatively invariant parameters such as binding and dissociation rates, more variable parameters which describe epigenetic states or potential crosstalk from other pathways can have strong effects on cell fate heterogeneity and need to be considered for decision-making.

Velocity, on the other hand, encompasses both position as well as system parameters. Even a two-dimensional velocity considering only p21 and Cdk2 can provide more insight than just p21 and Cdk2 levels as it contains information inherent to the entire regulatory network, including all interacting biochemical species, rate constants associated with these interactions, and epigenetically driven cell-to-cell variations in synthesis and degradation. In essence, velocity measured on a low-dimensional landscape aids in better specifying the overall cellular position on the more complex high-dimensional landscape underpinning cell behavior. Furthermore, this approach circumvents limitations in single-cell dynamic measurements imposed by the paucity of non-overlapping fluorophores required for distinguishing each species being observed. The increased information gained from such experimental velocity measurements also provides better insight into pre-existing cell states defined by parameter variations in highly complex systems^16,17,54^ and thereby improves predictability of cell behavior under perturbations. Following our observations in this study, we expect that analyses of trajectories in analogous cell-fate decisions would significantly benefit from the experimental measurement of system velocities, particularly under dynamic or transient inputs, providing insight and intuition into system features that are typically obscured in position-centric frameworks.

Finally, considering velocity as a modulator or predictor of cellular function can also improve strategies for altering system dynamics to engineer cell behavior. For instance, single-cell heterogeneity in cancer cell apoptosis under cisplatin was better explained by the rate of p53 accumulation rather than total p53 accumulated over time.^55,56^ Increasing the rate of accumulation (velocity) using a small molecule in these studies reduced the fraction of surviving cells after treatment independent of the total p53 accumulated under treatment. From a landscape perspective, these cells can be imagined as being given an initial cisplatin-driven impulse that directs them to a high-p53 apoptotic state while the landscape itself is biased towards a low-p53 survival state. Cells with high velocities from the initial impulse better resist the underlying landscape that favors a surviving state. This results in a velocity-dependent cell-fate decision in which faster cells undergo apoptosis while slower cells survive. Subsequently, using a small molecule to speed up slower cells results in more cell death. In addition to systems where velocity explicitly plays a role in cell decision-making, this phenomenon is likely to also arise under transient cues where the landscape dynamically changes in response to the presence and absence of the cue. For example, pharmacodynamic effects in a clinical context can lead to changes in the underlying energy landscape during and after drug administration. It would be informative to map the energy landscape under these scenarios and examine the effect of inertia on cell transitions in this context. This information could be further applied to formulate strategies based on exploiting inertia for favorable tumor responses.

In conclusion, this study highlights the role of velocity in determining cell fate on an energy landscape, using the quiescence-proliferation decision of breast cancer cells under hypoxia as a model system. This manifests as an inertial effect wherein the cells resist changes in their direction of movement even when the underlying landscape has changed to restrict movement in that direction (**Figure 4**). Interestingly, this velocity-driven inertia can even supersede the effects of initial position on the changed landscape, as seen in our study. This phenomenon is not limited to the cell cycle and is likely present in most non-equilibrium systems such as the circadian rhythm, p53, and NF-κB networks wherein external perturbations act on cells that are already “moving” on the landscape as well as systems where there is transience in the cue leading to reversible changes in the underlying landscape. Utilizing this additional information can provide valuable insight into the response of biological systems under perturbations, thereby enabling the design of effective strategies to modulate system response.

**Figure 4:**
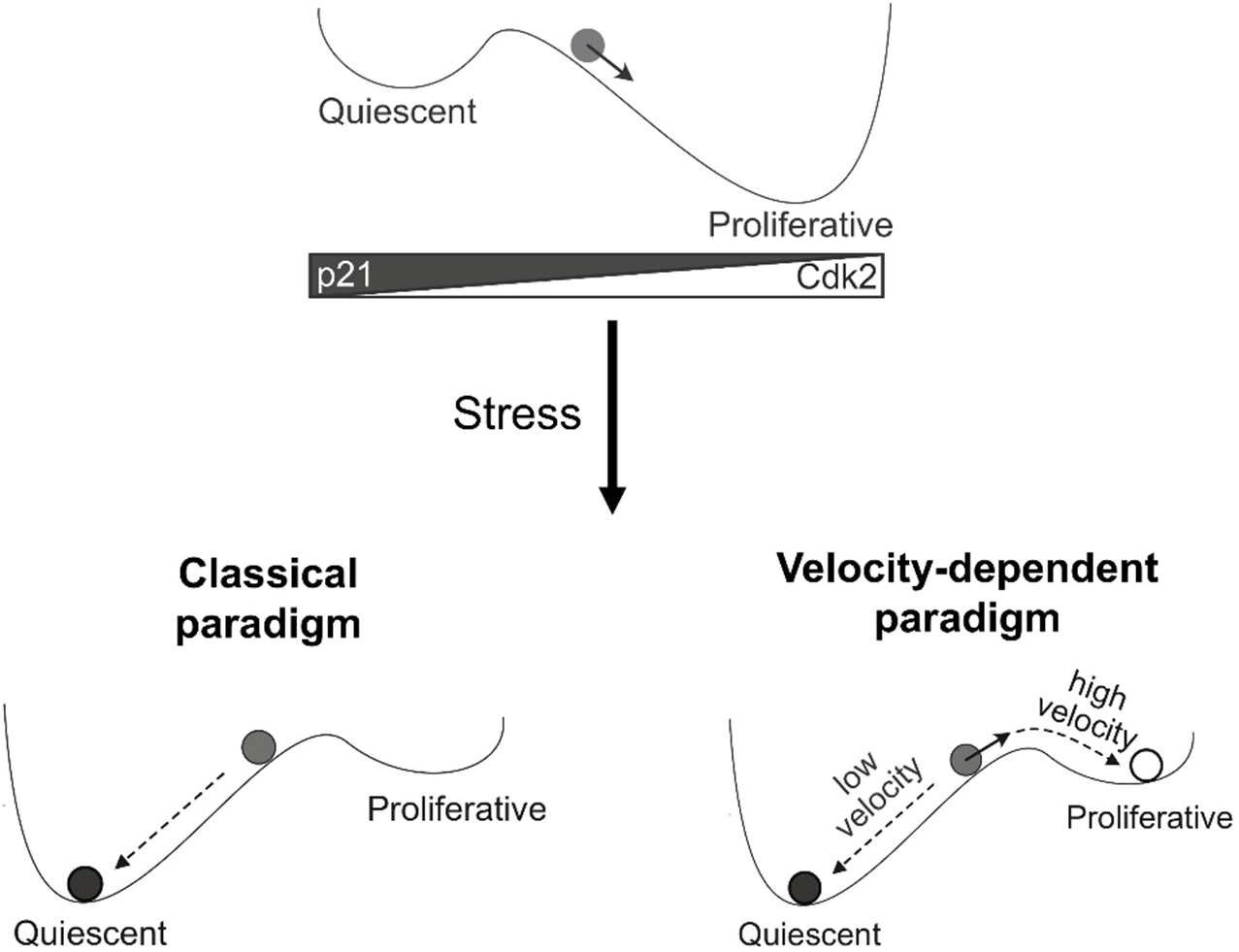
Cell decision-making on the p21-Cdk2 energy landscape under the classical position-only paradigm and the velocity-dependent paradigm. Cells are typically proliferative under normal conditions. However, under stress, the landscape becomes biased towards quiescence. In the classical paradigm, we would expect all cells in the indicated position to enter quiescence. However, we observe that fates are velocity-dependent, with high-velocity cells having sufficient inertia to overcome the change in landscape conditions and continue being proliferative.

## Supporting information

Code_Inertia_p21Cdk2.zip

## Acknowledgements

We thank Ryan Hanson for assistance with live-cell imaging and helpful discussions regarding image processing and analysis. This work was supported by funding from the University of Minnesota (S.M.A.) and the National Institutes of Health (R35GM136309; C.A.S.), as well as access to high-performance computing resources from the Minnesota Supercomputing Institute and FACS services from the University Flow Cytometry Resource at the University of Minnesota.

## Author Contributions

Conceptualization, H.V., S.M.A., C.A.S.; Methodology, H.V., E.B., S.M.A., C.A.S.; Software, H.V.; Formal analysis, H.V., C.B., E.B., S.M.A., C.A.S.; Investigation, H.V.; Resources, E.B., S.M.A., C.A.S.; Writing – Original Draft, H.V., S.M.A., C.A.S.; Writing – Review & Editing, H.V., E.B., S.M.A., C.A.S.; Supervision, E.B., S.M.A., C.A.S.; Funding acquisition, S.M.A, C.A.S.

## Materials and Methods

### Human cell lines and culture

MCF-7 and MDA-MB-231 human breast cancer cell lines (ATCC) were maintained in Dulbecco’s Modified Eagle’s Media (DMEM, 4500 mg/L glucose, MilliporeSigma) supplemented with 10% (v/v) fetal bovine serum (FBS; Thermo Fisher Scientific). MCF-7 cells tagged with mVenus fluorescent protein at the endogenous p21 locus and transfected with DHB-mCherry and H2B-mTurquoise (a kind gift from Dr. Sabrina L. Spencer^30^) were maintained in RPMI-1640 (MilliporeSigma) supplemented with 10% (v/v) FBS and 1% (v/v) penicillin-streptomycin (PenStrep 10,000 U/mL; Life Technologies). Due to the loss of DHB-mCherry and/or H2B-mTurquoise in subpopulations of these cells, fluorescence-activated cell sorting (FACS) was used to obtain cells that retained expression of both constructs.

### Live-cell microscopy and image analysis

For live-cell imaging, cells were plated on 6-well 1.5 glass-bottom dishes (Mattek) at a density of 1.25 x 10^5^ cells per well in transparent RPMI-1640 media (Life Technologies), lacking phenol red and riboflavin, supplemented with 10% (v/v) FBS and 1% (v/v) PenStrep. Cells were imaged with a Nikon Eclipse Ti2 inverted fluorescence microscope equipped with an automated stage (Prior) and a custom chamber to maintain 37 °C, 5% CO_2_, and high humidity. At each time point, images were collected for the brightfield, YFP, Texas Red, and CFP channels using a 20X CFI Plan Apochromat Lambda (NA=0.75) objective (Nikon) at 20% lamp intensity. Exposure time for each channel was set to be < 500 ms. The imaging protocol was paused briefly to add CoCl_2_ containing media to induce hypoxic conditions and resumed afterward. The ND2 format imaging data was exported as individual tiff files for each channel, time point, and stage position using Bio-Formats command line tools^57^. Cells were tracked semi-automatically in p53Cinema^58^ using the nuclear marker H2B-mTurquoise in the CFP channel. p21-mVenus expression for each cell trace was obtained using the getDatasetTraces_fillLineageInformation function while DHB-mCherry nuclear (DHB_nuc_) and cytoplasmic (DHB_cyt_) intensities were obtained by the getDatasetTraces_localSegmentation function, both built into p53Cinema. The traces were analyzed using MATLAB as described in the **Supplementary Information**. Briefly, the traces were background subtracted, smoothed, and clustered using dynamic time warping and hierarchical clustering. Scripts used are provided in **Code_Inertia_p21Cdk2.zip**.

### Cobalt chloride treatment to mimic hypoxia

CoCl_2_ (MilliporeSigma) was dissolved in type I Milli-Q water at a concentration of 10 mM and filtered using a sterile 0.45 µm polyethersulfone (PES) Steriflip 50 mL filter unit (MilliporeSigma). The resulting aqueous solution was directly added to the cell culture media to obtain a final concentration of 300 µM for MCF-7 cells and 100 µM for MDA-MB-231 cells, consistent with a prior study^28^. Treatment was initiated when the cells began exponential growth, as determined by growth curves (Day 2 and Day 3 for MDA-MB-231 and MCF-7 cells, respectively).

### Immunostaining

Cells in a 12-well plate were rinsed 3x with 1 mL DPBS, fixed with 4% paraformaldehyde for 15 min at room temperature, and washed 3x with 1 mL DPBS for 5 min each. Blocking and permeabilization were performed for 1 h in 1 mL blocking buffer - Dulbecco’s phosphate buffered saline (DPBS) containing 5% (v/v) normal goat serum (Thermo Fisher Scientific) and 0.3% (v/v) Triton X-100 (MilliporeSigma). Then, the cells were stained overnight at 4 °C with 500 µL solution of primary antibody diluted in antibody dilution buffer (DPBS containing 1% g/mL bovine serum albumin (MilliporeSigma) and 0.3% (v/v) Triton X-100). Antibodies and dilutions are provided in **Table S1**. The cells were then washed 3x with 1 mL DPBS for 5 min each followed by labeling for 1 h at room temperature with 500 µL of the corresponding secondary antibodies diluted in antibody dilution buffer. Cells were again washed 3x with 1 mL DPBS for 5 min each followed by nuclear labeling for 5 min with 1 mL 300 nM DAPI dissolved in DPBS (Thermo Fisher Scientific) and a 5-min DPBS wash. Finally, 1 mL DPBS was added to the cells before imaging. Samples were imaged using an EVOS FL Auto fluorescence microscope (Thermo Fisher Scientific). Quantification was performed with ImageJ software (National Institutes of Health). For imaging in other well formats, the volumes were scaled as per the typical culture media volume for each dish.

### Western blotting

For protein extraction, cells were washed twice with ice-cold PBS and lysed using RIPA buffer mixed with HALT Protease Inhibitor and HALT Phosphatase Inhibitor (Thermo Fisher Scientific). Whole-cell lysates were separated on an SDS-PAGE gradient gel (4–15%) and transferred to a polyvinylidene difluoride (PVDF) membrane (Bio-Rad). p21 and Cdk2 were detected in 15 μg of whole-cell lysates. β-actin was detected in 10 μg of whole-cell lysates. The membranes were blocked in 5% non-fat dry milk (Bio-Rad) in Tris-buffered saline containing 0.05% Tween-20 (TBS-T; MilliporeSigma) at room temperature for 1 h. Next, the membranes were incubated overnight at 4 °C with primary antibody diluted in 5% milk or 5% BSA in TBS-T (as per manufacturer’s protocol), followed by 1x 15-min wash and 2x 5-min washes with TBS-T. Antibodies and dilutions are provided in **Table S1**. The membrane was then incubated with its corresponding secondary antibody diluted in 5% milk in TBS-T at room temperature for 1 h, followed by TBS-T washes as with primary antibody. Protein expression was detected using SuperSignal West Pico Chemiluminescent substrate (Thermo Fisher Scientific) and a ChemiDoc Imager (Bio-Rad). Western blot quantification was performed by ImageLab (Bio-Rad). An equal-sized rectangular region was annotated around the detected protein band in each lane, and the integrated intensity within this box was calculated. This integrated intensity was background-corrected using a global background calculated using a region on the image not containing any protein bands or ladder. The background-corrected values were then normalized to β-actin expression, also calculated with the same processing workflow as described, from each sample to obtain the normalized protein expression values.

### Statistical analysis

Western blot data for each protein were analyzed using ANOVA followed by a multiple comparison test in GraphPad PRISM. Data obtained from time-lapse microscopy and single-cell tracking were analyzed using MATLAB. Comparisons of ranks were carried out using a non-parametric version of ANOVA, the Kruskal-Wallis test. Discrete distributions were compared using a custom MATLAB script implementing the Kolmogorov-Smirnov (KS) test where the statistical significance of the KS statistic was determined by comparison to a distribution of KS statistics calculated from 10^4^ shuffling iterations of the initial dataset. The statistical significance of same cluster vs. different cluster probabilities for sister cells was calculated by comparison with a distribution of probabilities calculated from 10^5^ shuffling iterations of the cluster identities within the original dataset.

### Computational methods

The ODE-based deterministic mathematical model^38^ was solved using the numerical stiff solver ode23s in MATLAB (Mathworks, Natick, MA). The stochastic version of the model was programmed and solved via the Gillespie algorithm^59^ in Julia^60^ using the ModelingToolkit.jl^61^ and DifferentialEquations.jl^62^ packages. For the purposes of landscape generation, we removed equations involving DNA synthesis, DNA damage induction and repair, as well as terms describing disassembly of DNA replication complexes post-S phase. Histograms, pseudo-energy landscapes and parameter phase spaces were visualized using MATLAB. Scripts used are provided in **Code_Inertia_p21Cdk2.zip**.

### Pseudo-energy landscape generation

Trajectories from the stochastic model were used to generate the pseudo-energy landscape as described previously.^34^ Briefly, the stochastic version of the model was run for multiple different initial values of total p21 and active Cdk2, each sampled from uniform distributions spanning the variable ranges of interest. The resulting probability histogram of the total p21 and active Cdk2 values from these trajectories was then converted into a pseudo-energy landscape using *U* = - ln(*P*) and visualized using the *surf* function in MATLAB. The number of trajectories (5000), discretization intervals (0.2 min), and simulation time (10,000 min) were determined by examining landscape convergence properties as described previously.^34^ Scripts used are provided in **Code_Inertia_p21Cdk2.zip**.

### Pseudo-energy landscape generation from experimental data

Analogous to the computational landscape methodology, we generated a two-dimensional histogram of mean p21 intensity and Cdk2 activity from time-lapse imaging data. We selected parts of the cell trajectories that were representative of cell fate and not primarily consisting of transitory cells that arise due to changes in the underlying landscape. As the cells were cultured under normoxia, they could be considered non-transitory and therefore, all traces prior to hypoxic treatment were used to generate the normoxic landscape. However, for the hypoxic landscape, we discarded cell trace values prior to 52 hours (∼1.5 times the average MCF-7 cell doubling time) of hypoxia to avoid overweighting the dynamics of the transition from the normoxic landscape to the hypoxic landscape. This two-dimensional probability distribution was converted to a potential energy landscape by taking its negative logarithm *U* = - ln(*P*) and all points with infinite energy due to zero probability were assigned the highest non-infinite energy. This resulting landscape was smoothed using a Gaussian-weighted filter (*smoothdata* function in MATLAB) to obtain the final energy landscape.

