## Supplementary Information for "Inertial effect of cell state velocity on the quiescence-proliferation fate decision in breast cancer"

**Contents**

Supplementary Methods: time-lapse microscopy and single-cell tracking; deterministic model simulations

Supplementary Table S1

Supplementary Figures S1 – S13

### Supplementary Methods

#### *Time-lapse microscopy and single-cell tracking*

Scripts used are provided in **Code_Inertia_p21Cdk2.zip.**

*Single-cell trace processing*

The individual cell traces obtained from p53Cinema were background corrected and smoothed for further analysis. p21-mVenus traces were background corrected by subtracting the minimum value of each individual trace from itself. For DHB-mCherry, the minimum of the cytoplasmic values or the nuclear values for each trace, whichever was lower, was subtracted from both the cytoplasmic and nuclear traces. Normalized Cdk2 activity (Cdk2a) is calculated as $DHB_{cyt}/(DHB_{cyt} + DHB_{nuc})$. In contrast to the ratio of cytoplasmic intensity to nuclear intensity ($DHB_{cyt}/DHB_{nuc}$), this scaled measure is more robust to temporal variations in sensor expression observed in our study (**Supplementary Figure 13**) while still being representative of Cdk2 activity. However, it must be noted that there appears to be a reduction in dynamic range in untreated cells.

*Data processing and clustering analysis*

Individual traces were smoothed using a moving-average algorithm in the *smoothdata* function in MATLAB. Dynamic time warping, *dtw* function in MATLAB, using a maximum warping distance of 5 data points (100 min in real world time) was then applied to calculate pair-wise dissimilarities between traces for both p21 and Cdk2. The Euclidean sum (sum of squares) of the p21 trace distance and Cdk2 trace distance was calculated for each pair of traces. These pair-wise distances were used to perform agglomerative hierarchical clustering (*linkage* function in MATLAB) with the 'ward' algorithm, previously shown to perform satisfactorily for single-cell traces[1]. The appropriate number of clusters was determined by calculating the Calinski-Harabasz index for different cluster numbers and picking the number with the highest value (*evalclusters* function in MATLAB); this corresponded to the number of clusters such that the ratio of inter-cluster variance to mean intra-cluster variance was maximized. In other words, on average, data was classified such that points within the same cluster had the highest similarity while being dissimilar to data points from any other cluster. In our dataset, the maximum was obtained for a choice of 4 clusters. However, one of the clusters consisted of very few cells compared to the other clusters. This cluster was considered as an outlier and discarded.

*Cell cycle classification*

Cells were classified into $G_0/G_1/S$, $S/G_2$, or M phases by the relative Cdk2 activity trend starting from the previous division up to the $CoCl_2$ addition. First, each trace was aligned to the last division before $CoCl_2$ was added to the culture. The traces were smoothed further using a Gaussian algorithm to preserve the general trend while more aggressively removing noisiness. If a trace had a higher value at $CoCl_2$ addition compared to immediately after division, it was assigned to the $S/G_2$ bin; if the value was lower, it was assigned to the $G_0/G_1/S$ bin. If there was a maximum in Cdk2 activity where Cdk2 initially rose and then eventually came back down, those cells were assigned to the M bin. In addition, if at least 50% of the trace had Cdk2 activity less than a threshold value of 0.38, it was classified as $G_0/G_1/S$. This threshold was determined by the central minima of the bimodal distribution of Cdk2 activities, representing the likely discriminant value between high and low Cdk2 activity states.

***Deterministic model simulations***

Scripts used are provided in **Code_Inertia_p21Cdk2.zip.**

*Phase space generation*

Phase spaces were generated using the parameter values specified in the figures. All other parameters as well as initial conditions (except for a quiescent initial state) were the same as those used by Heldt et al.[2] To obtain the quiescent initial state, the system was simulated using a low p53 degradation rate (0.01 times the basal kDeP53 value) with initial conditions from Heldt et al.[2] until the system reached steady state. This steady-state value was used as the initial condition for quiescent initial states in bistability calculations.

*Sensitivity analysis for S phase entry time*

A Latin-hypercube sampling scheme (*lhsdesign* function in MATLAB) was used to generate 1,000 parameter sets that vary over a 2-fold range above and below the original values. The system was then simulated for 5,000 min (~83 h, much longer than typical expected $G_1$ lengths in proliferative cells). The simulation was then classified as proliferative or quiescent based on whether DNA synthesis had occurred or not. In this model, DNA starts at 0 and reaches 1 when synthesis is completed. We used a threshold of 0.9 to evaluate whether DNA synthesis had occurred. For proliferative simulations alone, we calculated the time of S-phase entry which was defined as the time at which DNA reached a value of 0.1. We used a relatively high threshold to avoid confounding effects from the basal DNA synthesis that occurs in the model even prior to events classically associated with S-phase entry (rapid assembly of replication complexes and p21 degradation). In accordance with this criterion, the DNA synthesis rate constant kSyDna was held constant in all simulations as changes in this would have changed the time at which a threshold DNA constant would be reached directly and thereby would have obscured more biologically relevant effects. We then fitted a linear model using $\log_{10}$(S-phase entry time) as the output and $\log_2$(Parameters) scaled from 0 to 1 as the input (*fitlm* function in MATLAB). The coefficients of this model were used as a measure of the sensitivity of S-phase entry time to each parameter.

*Correlation of average total p21 and active Cdk2 with cell cycle velocity*

Average total p21 and average Cdk2 activity for each cell were measured *in vitro* over the entire cell cycle. However, the model does not cover the entire cell cycle; it does not loop back to $G_1$, but rather stays in $G_2$ indefinitely after S-phase exit. To have a more commensurate comparison, we considered the values of p21 and Cdk2 for 4 h after S-phase exit in the model, measured using DNA concentration thresholds as described in the previous section. This duration is the average eukaryotic $G_2$ phase length[3]. To qualitatively simulate the effects of biological heterogeneity, the E2f and p21 synthesis rates were varied from the basal values by -0.1 to 0.2 on the $\log_2$ scale and 0 to 1 on the $\log_{10}$ scale respectively. Additionally, each parameter pair was simulated 10 times where the initial values for CyclinD, total CyclinE:Cdk2, total CyclinA:Cdk2, and total p21 were sampled from a normal distribution centered around the values in Heldt et al.[2] with 20% coefficient of variation. Cell cycle velocity was considered the inverse of S-phase exit time. Quiescent cells were assigned a velocity of zero and their p21 and Cdk2 levels were averaged over the entire simulation length. Average levels of total p21 and active Cdk2 complexes were then measured and plotted as a function of cell cycle velocity.

**Table S1:** Antibodies used in this study with applications and dilutions

| Antibody | Manufacturer | Application & dilutions |
|---|---|---|
| Goat anti-rabbit Alexa Fluor 594 (Cat # A11012) | Thermo Fisher Scientific | Immunostaining (1:1000) |
| Goat anti-mouse Alexa Fluor 488 (Cat # A11001) | Thermo Fisher Scientific | Immunostaining (1:1000) |
| Rabbit anti-human Ki67 (D3B5) (Cat # 9129S) | Cell Signaling Technology | Immunostaining (1:1000) |
| Rabbit anti-human p21 primary monoclonal antibody (12D1) (Cat # 2947S) | Cell Signaling Technology | Immunostaining (1:1000), Western blotting (1:1000) |
| Mouse anti-human Cdk2 primary monoclonal antibody (D-12) (Cat # sc-6248) | Santa Cruz Biotechnology | Immunostaining (1:50), Western blotting (1:1000) |
| HRP (horseradish peroxidase)-conjugated β-actin (13E5) (Cat # 5125S) | Cell Signaling Technology | Western blotting (1:1000) |
| Goat anti-mouse polyclonal HRP-conjugated secondary antibody (Cat # 31430) | Thermo Fisher Scientific | Western blotting (1:6000) |
| Goat anti-rabbit polyclonal HRP-conjugated secondary antibody (Cat # 7074S) | Cell Signaling Technology | Western blotting (1:1000) |

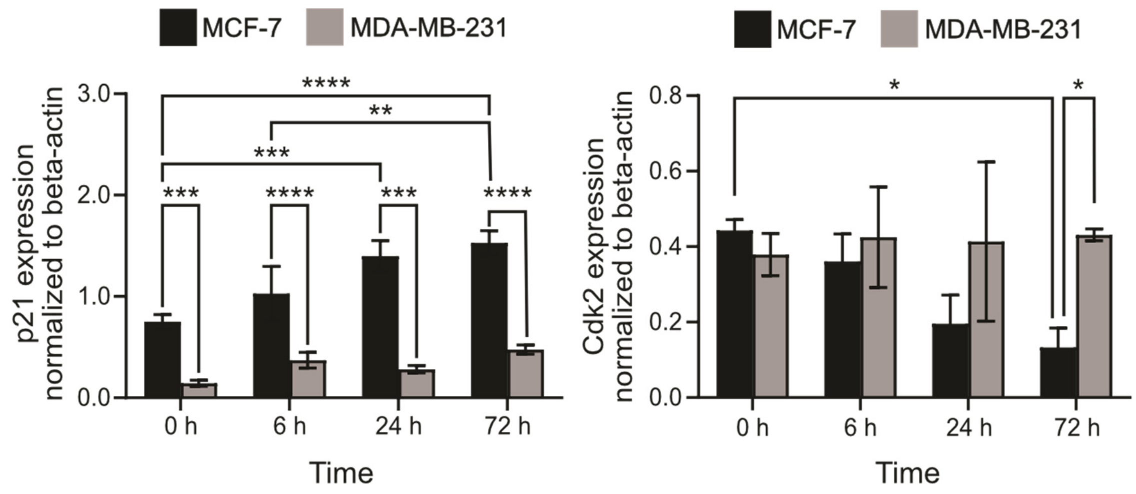


**Supplementary Figure 1. Quantification of p21 and Cdk2 western blots for MCF-7 and MDA-MB-231.** Bars represent mean and error bars represent standard deviation of p21 or Cdk2 expression in three biological replicates. (* $p < 0.05$; ** $p < 0.01$; *** $p < 0.001$; **** $p < 0.0001$)

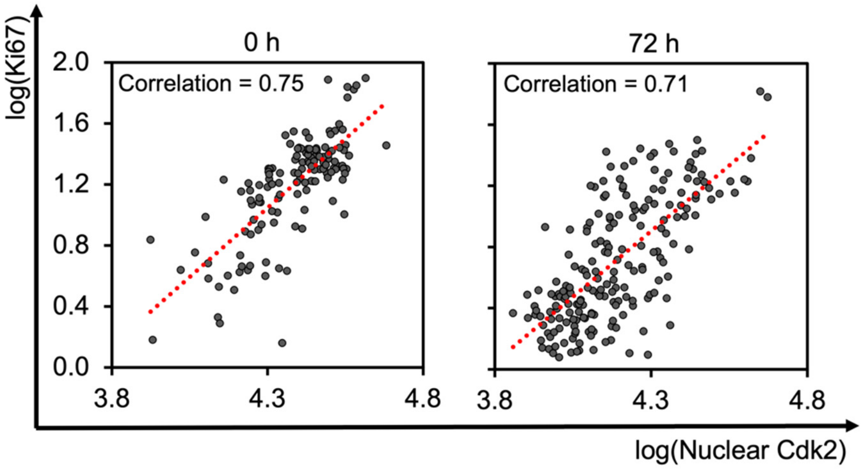


**Supplementary Figure 2. Single-cell expression of nuclear Cdk2 and Ki67.** The scatter plots show the logarithm of the integrated fluorescence intensities of Ki67 and Cdk2 within the nucleus of each cell as measured using immunofluorescence microscopy. Subheadings indicate the amount of time the sample was treated with $CoCl_2$. Red line represents a linear fit of the data, with the Pearson correlation coefficient of the data on the top left of the graph.

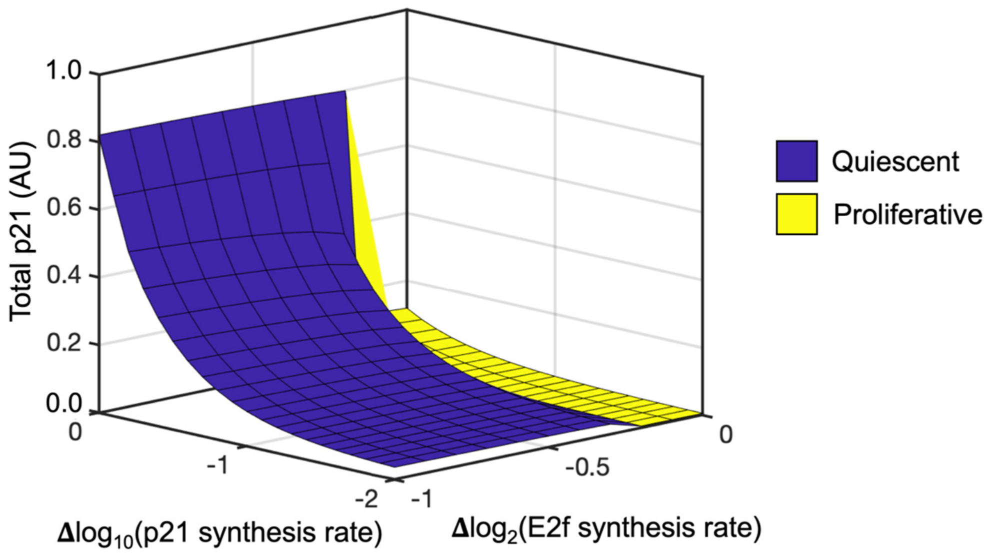


**Supplementary Figure 3: Total p21 at steady state as a function of the relative change in p21 synthesis rate and E2f synthesis rate.** Relative change is calculated by dividing the current parameter value by the basal value and taking its logarithm. Blue and yellow regions correspond to quiescence and proliferation, respectively. The region near $10^{-2}$ fold change in p21 synthesis rate and $2^{-1}$ fold change in E2f synthesis rate represents a quiescent steady state governed by low mitogenic signaling as opposed to high p21 upregulation. This is representative of a serum-starved state reported by Barr et al.[4]

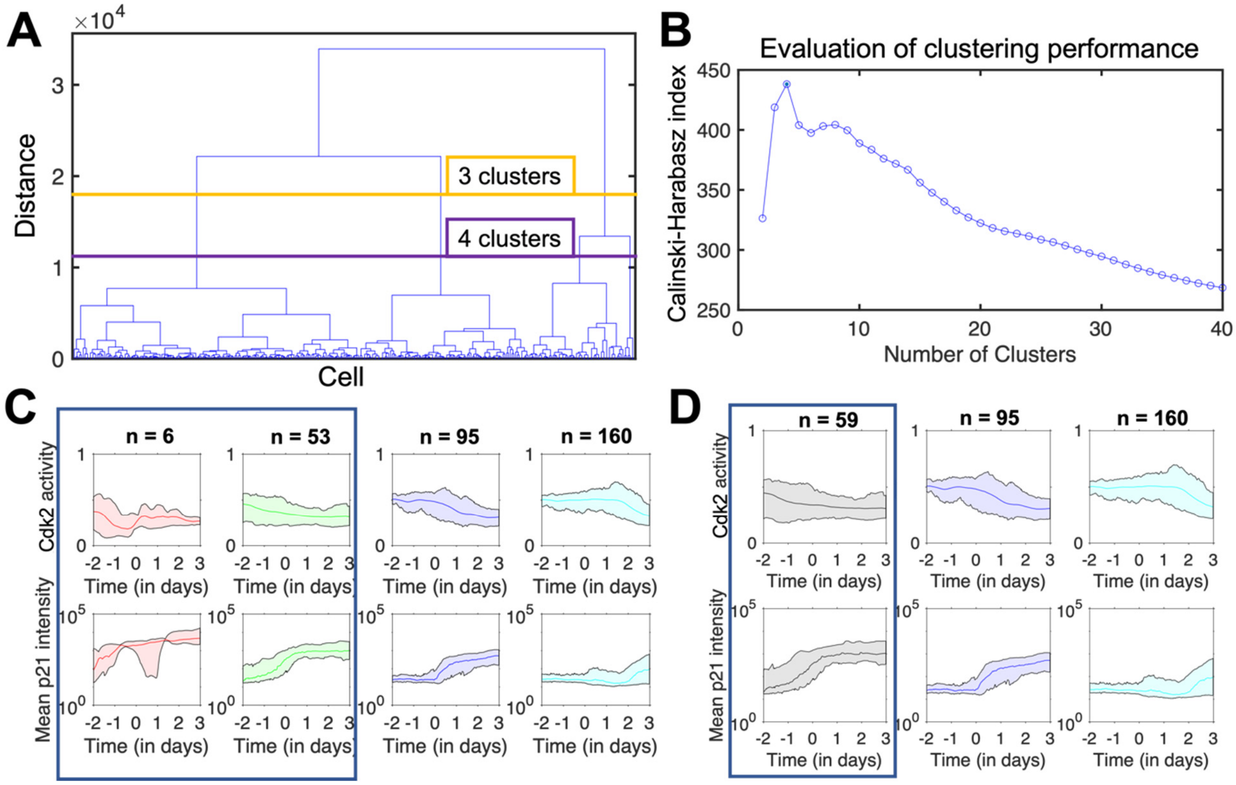


**Supplementary Figure 4: Unsupervised clustering of p21-Cdk2 cell traces. (A)** Dendrogram showing hierarchical clustering of cell traces. Each element being clustered represents one cell; the yellow and purple horizontal bars represent cutoff distances for coarse-grained classification into 3 and 4 clusters, respectively. **(B)** Plot of the Calinski-Harabasz values representing the ratio of inter-cluster to intra-cluster variances as a function of cluster number. The optimal number of clusters is 4. **(C, D)** Time-course plots showing Cdk2 activity and mean p21 intensity for 4 clusters (C) and 3 clusters (D). Solid lines in color represent cluster means and the surrounding areas in color bounded by black lines represent the range of $10^{th}$ and $90^{th}$ percentiles. Classifying all of the data into 3 clusters instead of 4 (i.e., not discarding one of the clusters as an outlier due to a low number of cells) does not significantly change the overall system behavior in the combined cluster.

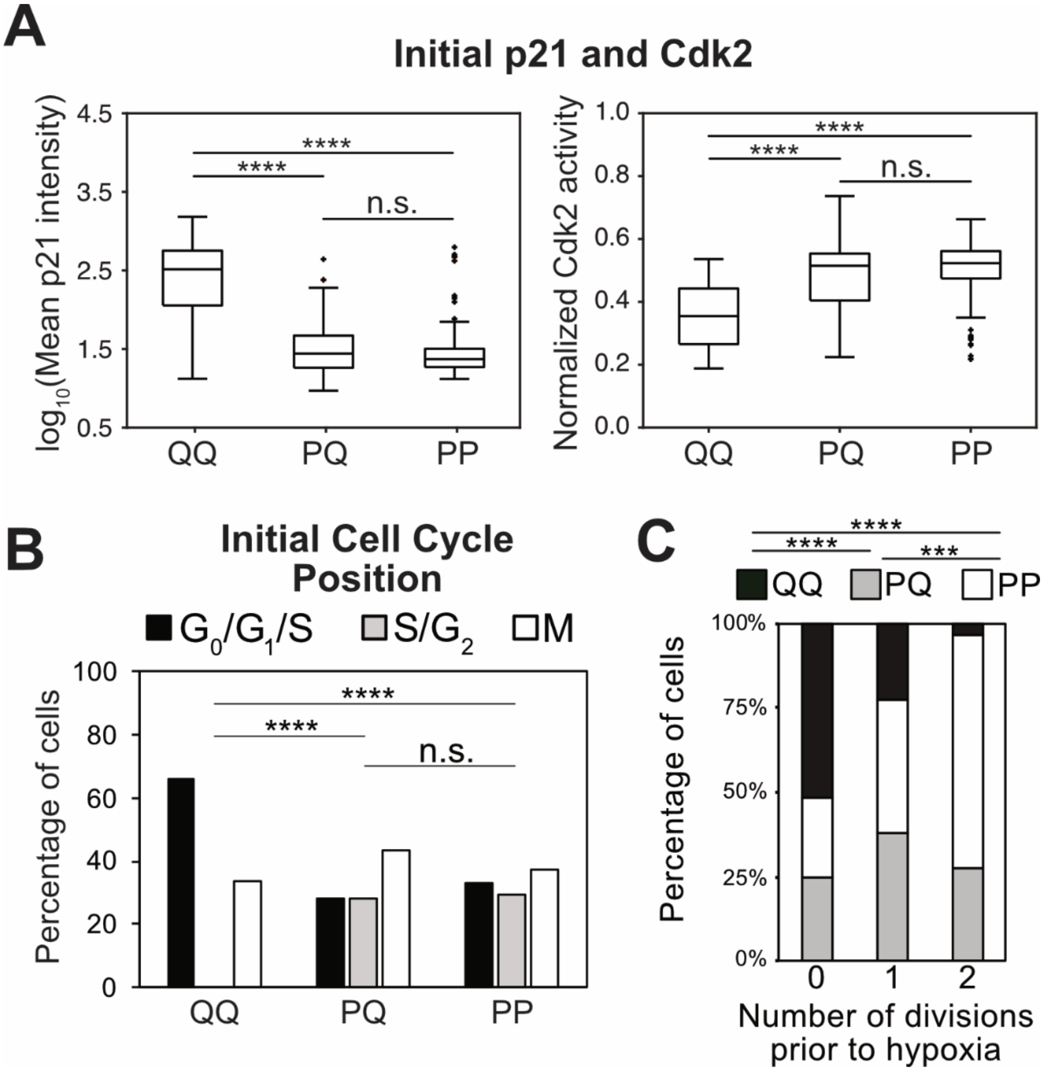


**Supplementary Figure 5: Effect of initial position and velocity on cell fate when all daughter cells are considered independent of each other. (A, B)** Box plots of p21 and Cdk2 expression **(A)** and cell-cycle position distributions **(B)** classified according to cluster identity showing differences between QQ and PQ/PP but not between PQ and PP. **(C)** Distribution of cell-fate cluster identity depending on number of divisions prior to hypoxia showing differences among all three clusters. (* $p < 0.05$; ** $p < 0.01$; *** $p < 0.001$; **** $p < 0.0001$) The conclusions drawn from these results support those in the main text. After hypoxia, < 5% of all cells (12 out of 255) had divergent lineages with daughters belonging to different clusters, while 39 out of 255 cells divided with daughters following the same lineage, implying that considering each cell as an independent individual effectively contributes significant double counting that can skew statistical analyses.

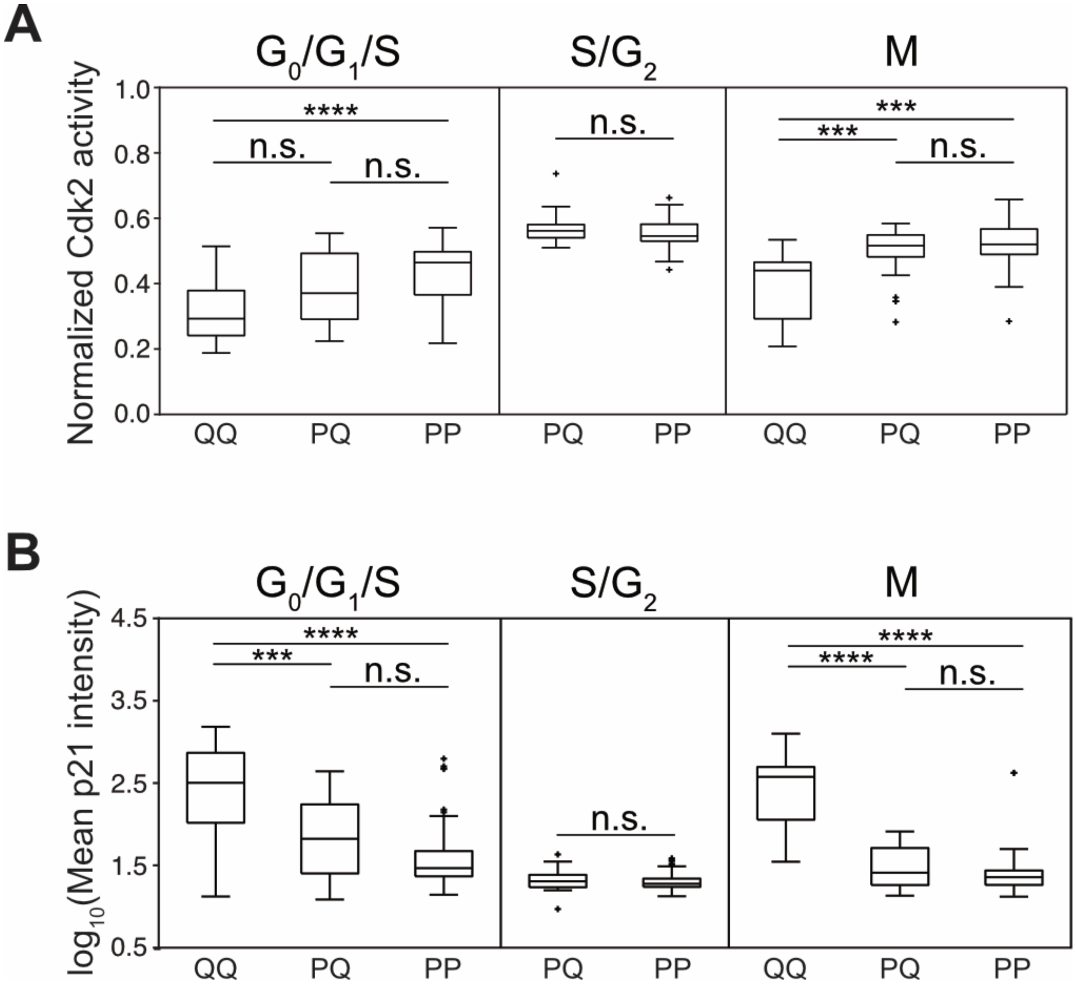


**Supplementary Figure 6: p21 and Cdk2 expression classified according to approximate cell cycle phase and then cluster identity.** Taken together, these plots show that there is a difference in position determined together by p21 and Cdk2 levels between clusters QQ and PQ/PP, while there are no differences between clusters PQ and PP across all phases of the cell cycle. (* $p < 0.05$; ** $p < 0.01$; *** $p < 0.001$; **** $p < 0.0001$)

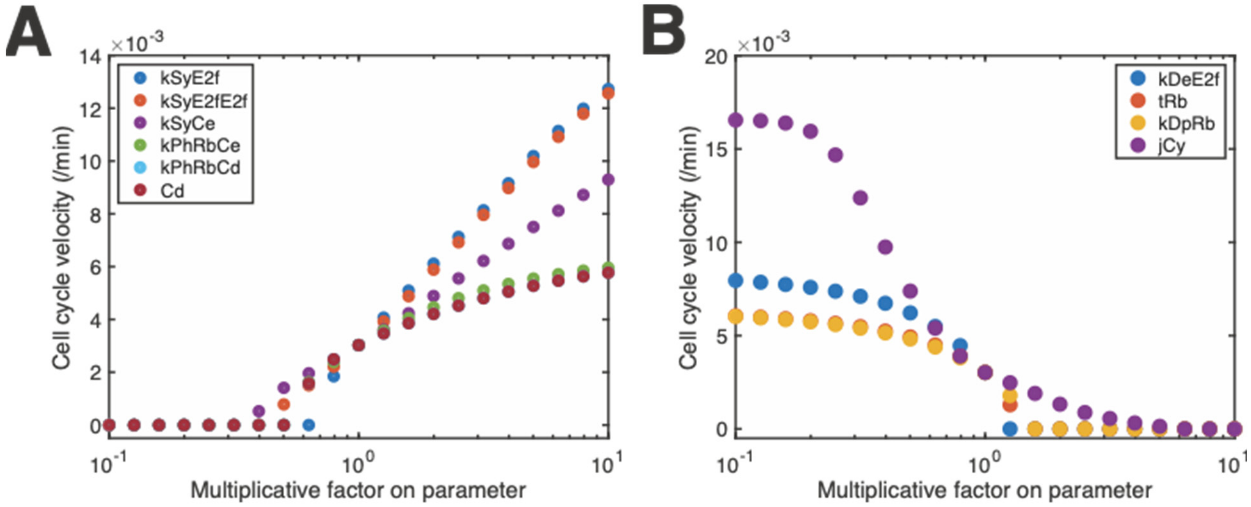


**Supplementary Figure 7: Effect of change in parameters on cell cycle velocity for parameters with (A) positive correlation and (B) negative correlation to velocity.** The change in parameter is measured as the multiplicative factor applied to the parameter from the range 0.1 to 10. **(A)** Changes in cell cycle velocity as a function of kSyE2f (basal E2f synthesis rate), kSyE2fE2f (E2f-dependent E2f synthesis rate), kSyCe (CyclinE synthesis rate), kPhRbCe (CyclinE-dependent Rb phosphorylation rate), kPhRbCd (CyclinD-dependent Rb phosphorylation rate), and Cd (total CyclinD). **(B)** Changes in cell cycle velocity as a function of kDeE2f (E2f degradation rate), tRb (total retinoblastoma (Rb)), kDpRb (Rb dephosphorylation rate), and jCy (Cyclin:Cdk2 complex threshold required for start of DNA synthesis).

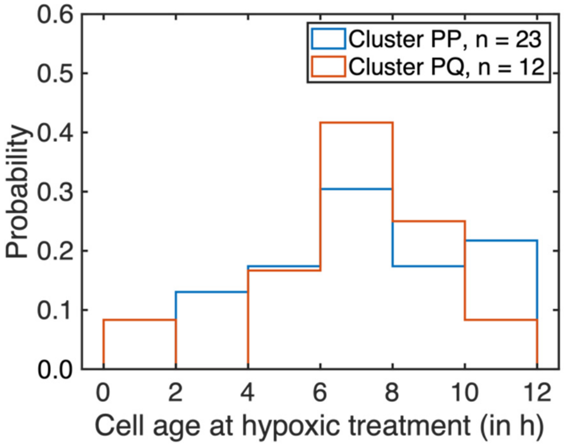


**Supplementary Figure 8: Probability distribution of the time after birth (mitosis) with respect to the time of treatment.** Blue and orange bars represent clusters PQ and PP respectively. A discretized version of the Kolmogorov-Smirnov test detected no statistically significant difference between the two distributions.

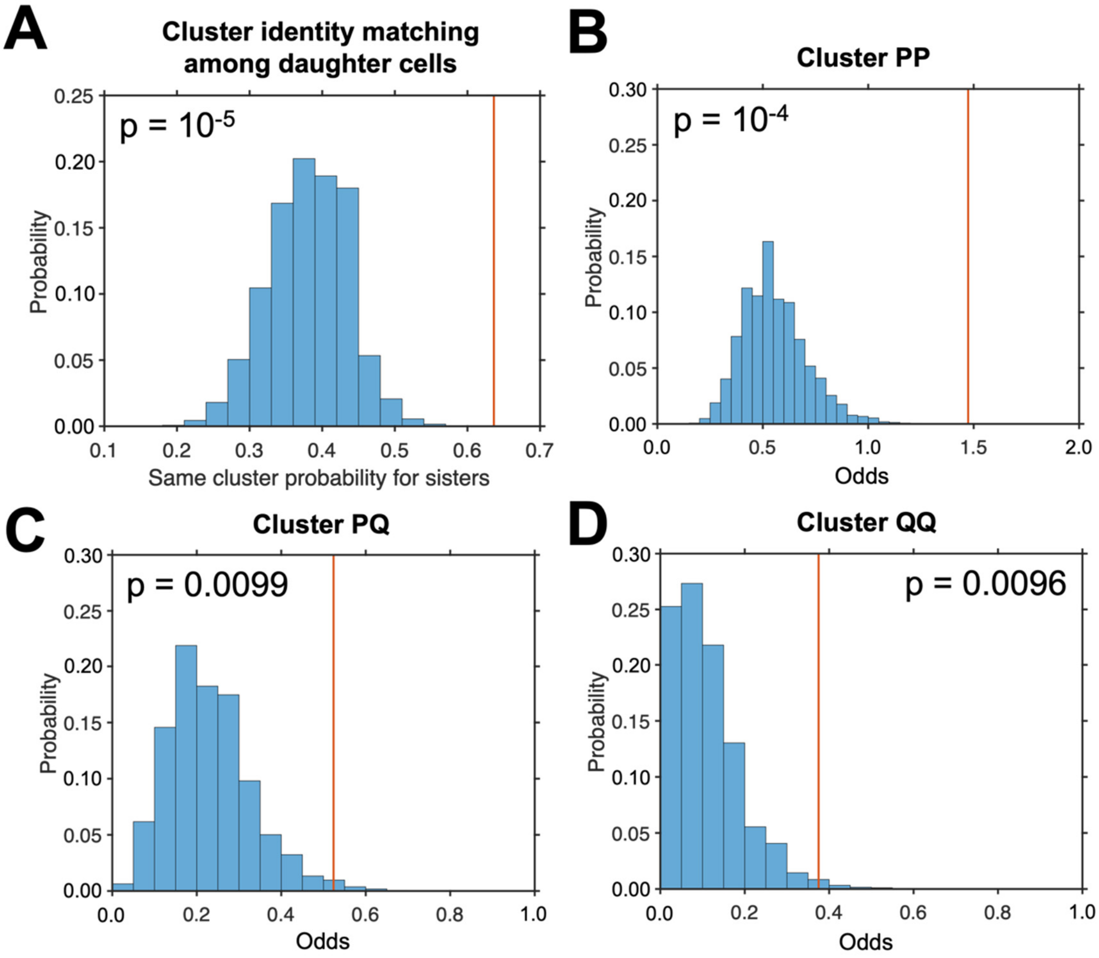


**Supplementary Figure 9: Statistical significance of inheritance probabilities. (A)** The distribution of the probability values of sister cells being in the same cluster obtained by randomizing the cluster identity of all cells (blue) compared to the calculated value for this dataset (orange line). **(B-D)** The distributions of the odds of both sister cells being in each given cluster versus being in different clusters obtained by cluster identity randomization (blue) compared to the calculated values for this dataset (orange lines).

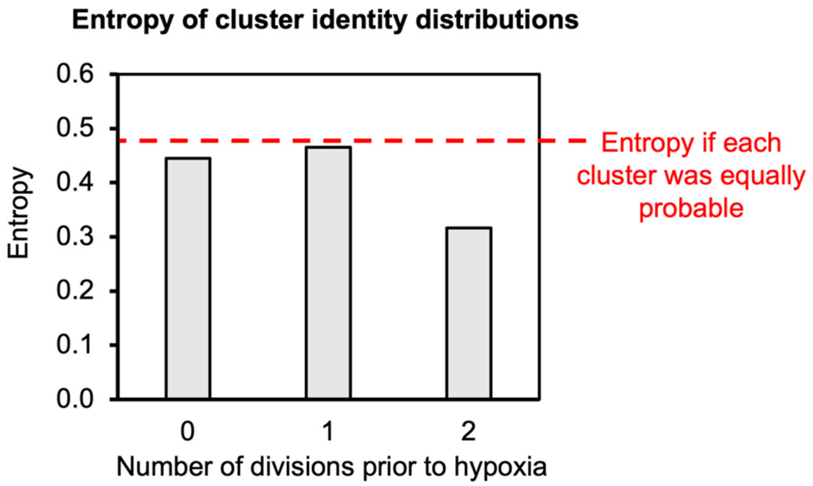


**Supplementary Figure 10: Entropy of cluster identity distributions as a function of number of divisions prior to hypoxia.** Entropy was calculated for cells that divided 0, 1, or 2 times prior to hypoxia using $E = -\sum p_i \cdot ln(p_i)$, where $i$ is the cluster identity QQ, PQ, or PP. Dashed red line represents the entropy of a uniform distribution with three bins, wherein each cluster identity is equally probable.

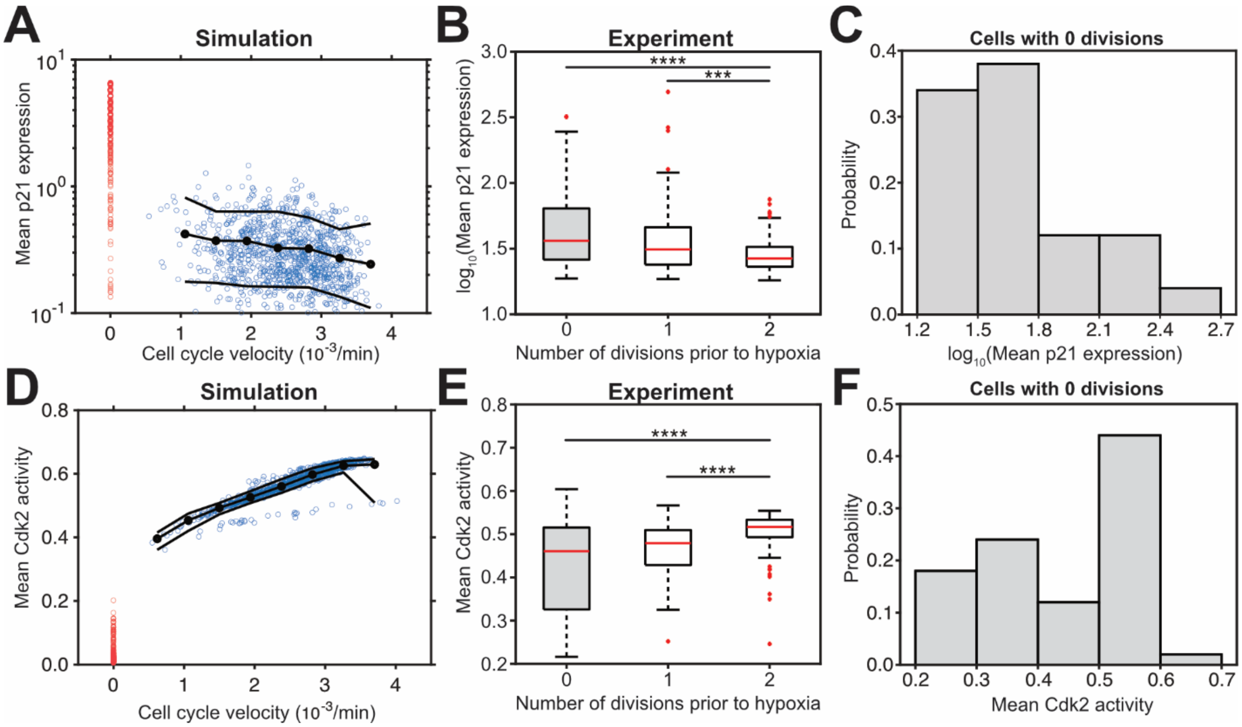


**Supplementary Figure 11: The effect of parameter-dependent cell cycle velocity on mean p21 expression and mean Cdk2 activity. (A)** Scatter plot of simulated mean p21 expression as a function of cell cycle velocity. Red and blue circles represent quiescent and proliferative cells, respectively. Line with black circles represents the mean of the blue circles at different cell cycle velocities, and solid black lines represent the standard deviations from the mean. **(B)** Box plots of the experimentally measured mean p21 expression as a function of the number of divisions prior to hypoxia with statistical testing on mean rank differences carried out using the Kruskal-Wallis test followed by a multiple comparison analysis. **(C)** Distribution of the mean p21 values of cells with 0 divisions prior to hypoxia (box plot in gray in panel B) showing high variability that likely underlies the lack of statistically significant difference between 0 and 1 divisions. **(D, E, and F)** Analogous plots for mean Cdk2 activity. (***, $p < 10^{-3}$ and ****, $p < 10^{-4}$)

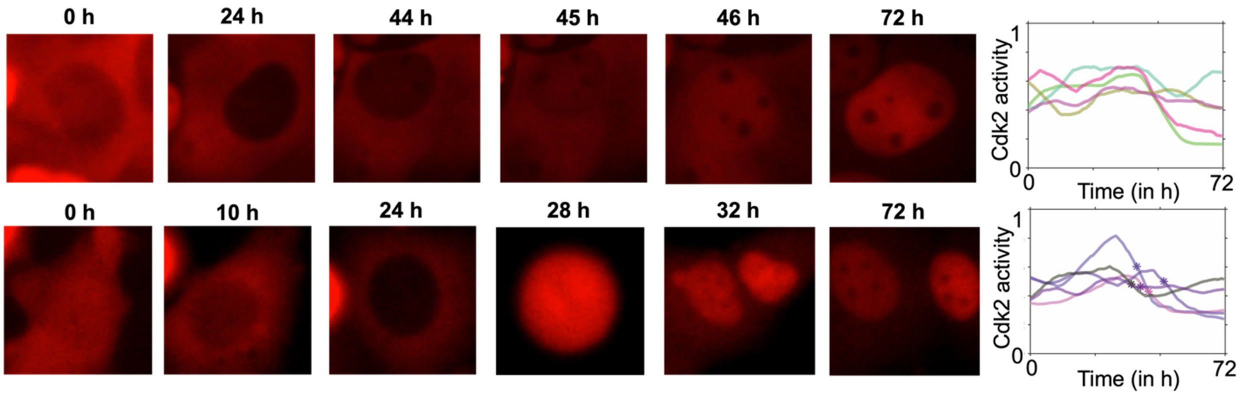


**Supplementary Figure 12: Cells that initially remain proliferative under hypoxia treatment eventually enter quiescence.** Representative images and Cdk2 activity traces for cells that continue to proliferate immediately after hypoxia but eventually enter a low Cdk2 state with (top) or without (bottom) cell division. Cell divisions in the bottom panel are noted by stars on each trace.

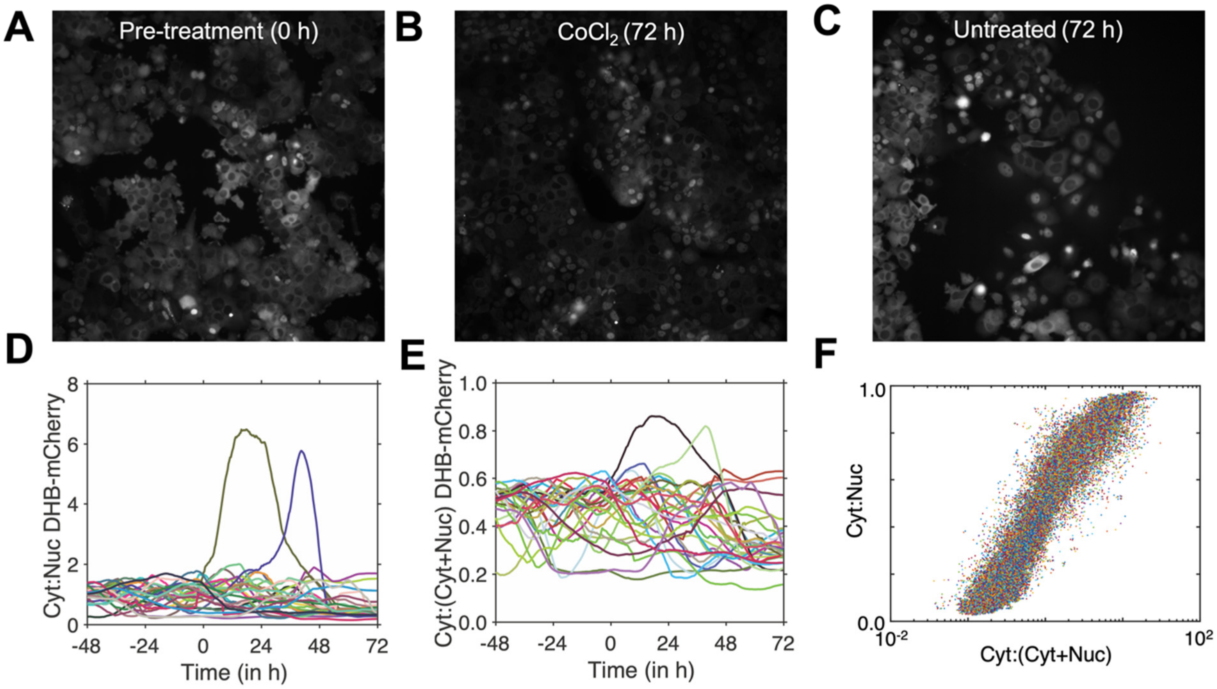


**Supplementary Figure 13: Use of a normalized fluorescence ratio to measure Cdk2 activity. (A)** DHB-mCherry expression at 0 h (pre-treatment). **(B)** DHB-mCherry expression after 72 h of cobalt chloride treatment with the same field of view as the 0 h image showing a significant change in sensor expression, likely due to dormancy induction. **(C)** DHB-mCherry expression after 72 h in an untreated well; these cells show comparable fluorescence to the 0 h time point. **(D)** Time series of Cdk2 activity in a subset of the total dataset as measured by the ratio of cytoplasmic to nuclear fluorescence, showing traces that have large aberrant values post-cobalt chloride treatment compared to pre-treatment, likely due to a drop in expression of the Cdk2 sensor leading to extremely low nuclear fluorescence values. **(E)** Time series of data in **(D)** where Cdk2 activity is measured by the ratio of cytoplasmic fluorescence to the sum of the nuclear and cytoplasmic fluorescence showing a more constrained range of output values (0 to 1). **(F)** Scatter plot of the cytoplasmic:nuclear ratio and the normalized ratio showing a strong correlation in activities.
