## Supplementary material for "Inertial effect of cell state velocity on the quiescence-proliferation fate decision in breast cancer": Code_Inertia_p21Cdk2.zip: AnalysisScript_p21Cdk2Traces.m

```
%% This code is for analysis of images using p53Cinema (Reyes et. al.)

%%
load('p21Cdk2Measurements_Hypoxia.mat')


%% CDK2 processing

CDK2_filt = CDK2;
CDK2_nuc_filt = CDK2_nuc;
CDK2_cyt_filt = CDK2_cyt;

%p53Cinema assigns unreadable data points to -1. For data processing, this
%is assigned as NaN
CDK2_nuc_filt(CDK2_nuc_filt < 0) = NaN;
CDK2_cyt_filt(CDK2_cyt_filt < 0) = NaN;
CDK2_filt(CDK2_filt < 0) = NaN;

%Background correction
%The minimum intensity in cytoplasmic and nuclear DHB-mCherry intensity for
%each cell is taken and then subtracted from both nuclear and cytoplasmic
%matrices. This improves the dynamic range of the normalized ratio
CDK2_bg = min(min(CDK2_nuc_filt,CDK2_cyt_filt)');

CDK2_bg_vec = repmat(CDK2_bg,size(CDK2,2),1)';

CDK2_nuc_filt_corr = CDK2_nuc_filt - CDK2_bg_vec;
CDK2_cyt_filt_corr = CDK2_cyt_filt - CDK2_bg_vec;


%Normalization of DHB-mCherry fluorescence (Ref. Materials and Methods)
CDK2_norm = CDK2_cyt_filt_corr./(CDK2_cyt_filt_corr +  CDK2_nuc_filt_corr);


%Smoothing of data using a moving mean
CDK2_smooth = (smoothdata(CDK2_norm','movmean'));

%Filling missing data by taking the nearest neighbor with a preference to
%the previous value. Avoids issues from interpolation of NaN and a number
%and accounts for cases where there are multiple NaNs in a row.
CDK2_smooth = fillmissing(CDK2_smooth,'previous',2);
CDK2_smooth = fillmissing(CDK2_smooth,'next',2);


%Defining the time vector for our experiment. Readings are taken once
%every 20 minutes with hypoxic treatment starting at 48 hours from start of
%the experiment. Induction of hypoxia is designated as 0 hours.
time_vec = (0:359)/3 - 48;

%Plotting individual cell traces
figure
for kk = 1:size(CDK2_smooth,2)
plot(time_vec,CDK2_smooth(:,kk),'Color',[rand rand rand 0.25])
hold on
end
hold off
xlabel('Time (in h)')
ylabel('Normalized Cdk2 activity')
set(gca,'TickDir','out')
set(gca,'FontName','Arial')
set(gca,'LineWidth',2)
set(gca,'FontSize', 24)
set(gca,'XTick', (-48:24:72))
set(gca,'YTick', (0:0.2:1))
ylim([0 1])
xlim([-48 72])


%% p21 processing

%Reading p21 intensities and setting unreadable values as NaN, similar to
%Cdk2
p21_s_filt = p21_sampled(:,:);
p21_s_filt(p21_s_filt<0) = NaN;


%Background correction similar to Cdk2 on a per cell basis
p21_bg = (repmat(min(p21_s_filt'),size(p21_sampled,2),1));

p21_s_filt = p21_s_filt' - p21_bg;


%Smoothing on lognormalized data, since the original intensities have a
%large range
p21_smooth = log10(smoothdata(p21_s_filt,'movmean'));
%Removing numbers that are zero intensity, and thus -Inf in log terms
p21_smooth(abs(p21_smooth)==Inf)=NaN;

%Filling missing data, similar to Cdk2
p21_smooth = (fillmissing(p21_smooth,'previous',2));
p21_smooth = (fillmissing((p21_smooth),'next',2));

%Time period, similar to Cdk2
time_vec = (0:359)/3-48;

figure
for kk = 1:size(p21_smooth,2)
plot(time_vec,p21_smooth(:,kk),'Color',[rand rand rand 0.25])
hold on
end
hold off
xlabel('Time (in h)')
ylabel('log10(Mean p21 intensity)')
set(gca,'TickDir','out')
set(gca,'FontName','Arial')
set(gca,'LineWidth',2)
set(gca,'FontSize', 24)
set(gca,'XTick', (-48:24:72))
set(gca,'YTick', (0.5:1:4.5))
ylim([0.5 4.5])
xlim([-48 72])


%% Dynamic time warping (DTW)-based distance calculations between each cell and dendrogram generation
rng(8.0) %Calling randomization to help with between run uniformity for data

n_cells = size(CDK2_smooth,2);

dist_mat = zeros(n_cells);


for i=1:(n_cells-1)
    for j=(i+1):n_cells
        %Calculate euclidean sum of between cell p21 and Cdk2 distances
dist = sqrt(dtw((CDK2_smooth(:,i)),(CDK2_smooth(:,j)),5)^2 + (dtw((p21_smooth(:,i)),(p21_smooth(:,j)),5))^2);

dist_mat(i,j) = dist;
dist_mat(j,i) = dist_mat(i,j); %Utilizes symmetry of the distance matrix
    end
end


Z = linkage((dist_mat),'ward'); %Generate hierarchical tree


leafOrder = optimalleaforder(Z,dist_mat);
ndivs_percell=sum(cell_divisions(:,1:146),2);
figure
dendrogram(Z,0,'reorder',leafOrder)


%% Code to evaluate clustering efficiency
eva = evalclusters(dist_mat, 'linkage','CalinskiHarabasz','Klist',1:40);
plot(eva)

%% Actual clustering and plotting
%From CH index we know it is 4 clusters.
max_clust = 4;

T = cluster(Z,'maxclust',max_clust);
time_vec = (0:359)/3 - 48;

for i = 1:max_clust
    subplot(2,max_clust,i)
    color_vec=[rand*1 rand*1 rand 0.2];
    l1 = plot(time_vec, (CDK2_smooth(:,T==i)),'Color',color_vec);
    pbaspect([1.5 1 1])
   ylim([0 1])
   xlim([-48 72])
    set(gca,'FontName','Arial')
    set(gca,'FontSize',16)
    xlabel('Time (in h)')
    xlim([min(time_vec) max(time_vec)])
    if (i==1)
    ylabel('CDK2 activity')
    end

    subplot(2,max_clust,i+max_clust)
    k2 = plot(time_vec,10.^p21_smooth(:,T==i),'Color',color_vec);
    pbaspect([1.5 1 1])    
    set(gca,'FontName','Arial')
    set(gca,'FontSize',16)
    xlabel('Time (in h)')
    if i == 1
    ylabel('Mean p21 intensity')
    end
    set(gca,'YScale','log')
    xlim([min(time_vec) max(time_vec)])
end


color_vec = {'red', 'green', 'blue', 'cyan', 'magenta', 'black','yellow'};

figure
j_vec = 1:max_clust;
for i = 1:max_clust
    
    j = j_vec(i);
        subplot(2,max_clust,j)
    meanCDK2vals = mean(CDK2_smooth(:,T==i),2);
    CDK2ranges = 0.5*iqr(CDK2_smooth(:,T==i),2);
    CDK2_75 = prctile(CDK2_smooth(:,T==i),90,2);
    CDK2_25 = prctile(CDK2_smooth(:,T==i),10,2);
    plot(time_vec, meanCDK2vals,'Color',color_vec{j})
    hold on
    patch([time_vec, flip(time_vec)], [CDK2_25; flip(CDK2_75)],'black','FaceColor',color_vec{j},'FaceAlpha', 0.1)

    xlim([min(time_vec) max(time_vec)])
    ylim([0 1])
    xlim([-48 72])
    set(gca,'XTick',(-48:24:72))
    set(gca,'XTickLabel',{'-2','-1','0','1','2','3'})
    set(gca,'TickDir','out')
    set(gca,'FontName','Arial')
    set(gca,'FontSize',16)
    xlabel('Time (in days)')
    if (j==1)
    ylabel('CDK2 activity')
    end
    subplot(2,max_clust,j+max_clust)
    meanp21vals = median((p21_smooth(:,T==i)),2);
    p21ranges = 0.5*iqr(log10(p21_smooth(:,T==i)),2);
    p21_75 = prctile(p21_smooth(:,T==i),90,2);
    p21_25 = prctile(p21_smooth(:,T==i),10,2);
    plot(time_vec, 10.^meanp21vals,'Color',color_vec{j})
    hold on
    patch([time_vec, flip(time_vec)], 10.^[p21_75; flip(p21_25)],'black','FaceColor',color_vec{j},'FaceAlpha', 0.1)
    set(gca,'FontName','Arial')
    set(gca,'FontSize',16)
    xlabel('Time (in days)')
    if j==1
    ylabel('Mean p21 intensity')
    end
    set(gca,'YScale','log')
    xlim([-48 72])
     set(gca,'XTick',(-48:24:72))
    set(gca,'XTickLabel',{'-2','-1','0','1','2','3'})
    set(gca,'TickDir','out')

   ylim(10.^[0 5])
end

%the following is for filtering out the outlying cluster. DO NOT USE THIS
%IF YOU ARE PERFORMING SISTER ANALYSIS as the way it is set it, it requires
%the presence of ALL cells in the lineagetree
% p21_smooth(:,T==1) = [];
% CDK2_smooth(:,T==1) = [];
% ndivs_percell(T==1) = [];
% lineagetree(T==1,:) = [];
% cell_divisions(T==1,:) = [];
% 
% T(T==1) = [];
% 
% T = T-1;

%% Generating trajectory-based landscapes and plotting cluster paths

%Inputs below are for normoxic landscape
% Input1 = CDK2_smooth(1:145,:);
% Input2 = p21_smooth(1:145,:);

%Inputs below are for hypoxic landscape
Input1 = CDK2_smooth(300:360,:);
Input2 = p21_smooth(300:360,:);

%Setting boundaries based on what our phase space of interest is. x is
%Cdk2, y is log10(p21)
xedges = 0.1:(0.9-0.1)/49:0.9;
yedges = 0.5:(4.0-0.5)/49:4.0;

%Generating the histogram based on trajectories
[h2] = histcounts2(Input1, Input2,xedges,yedges,'Normalization','Probability');

%Convert to -log for changing probability to energy landscape
U_landscape = -log(h2);

%Set singularities due to log transformation of zero probabilities to be
%the maximum potential observed on the landscape
U_landscape(U_landscape==Inf)=max(max(U_landscape(U_landscape~=Inf)));

xcenters = (xedges(1:end-1) + xedges(2:end))/2;
ycenters = (yedges(1:end-1) + yedges(2:end))/2;

%Plot the landscape
figure
surf(xcenters, ycenters, smoothdata(smoothdata(U_landscape,'gaussian',10)','gaussian',10),'EdgeAlpha',0)%surf(U_landscape)
hold on
colormap((jet))

% The section below plots the paths for each cluster

% %Use the jet colormap if you only want the landscapes. Use gray when you
% %also want paths that are plotted below also. 
% colormap(gray)
% time_range = 145:288; 
% Cluster_Identity = 1; %This the cluster whose path you want to plot
% for k = Cluster_Identity
%     meanCDK2vals = mean(CDK2_smooth(time_range,T==k),2);
%     meanp21vals = mean(p21_smooth(time_range,T==k),2);
%     CDK2_75 = prctile(CDK2_smooth(time_range,T==k),90,2);
%     CDK2_25 = prctile(CDK2_smooth(time_range,T==k),10,2);
%     p21_75 = prctile(p21_smooth(time_range,T==k),90,2);
%     p21_25 = prctile(p21_smooth(time_range,T==k),10,2);
%     CDK2bound = [CDK2_75; CDK2_25; CDK2_75; CDK2_25];
% p21bound = [p21_75; p21_25; p21_25; p21_75];
% zz = boundary(CDK2bound,p21bound,0.9);
% 
% fill3(CDK2bound(zz),p21bound(zz), 30*ones(size(CDK2bound(zz))),'k','FaceAlpha',0.5)
% 
% colormap_vec = jet(length(time_range));
% 
% for nn = 1:length(meanCDK2vals)-2
% p =  plot3(meanCDK2vals(nn:nn+2),meanp21vals(nn:nn+2), [50 50 50],'LineWidth',10,'Color',colormap_vec(nn,:));
% hold on
% end
% 
% hold on
% end
view(2)
ylim([0.5 4])
xlim([0.1 0.7])
set(gca,'XTick',[0.1 0.3 0.5 0.7])
set(gca,'XTickLabels',{'0.1','0.3','0.5','0.7'})
set(gca,'YTickLabels',{'0.5','1.0','1.5','2.0','2.5','3.0','3.5','4.0'})
set(gca,'Layer','top')
grid off
set(gca,'LineWidth',2)
set(gca,'FontSize',22)
set(gca,'Fontname','Arial')
set(gca,'TickDir','out')
pbaspect([1.1 1 1])
%
box on


%% Alignment to cell division
tmax = 48;
indmin = 1;
indmax = 145;
cell_divisions_norm = cell_divisions(:,indmin:indmax)';
CDK2_normoxia = CDK2_smooth(indmin:indmax,:);
p21_normoxia = p21_smooth(indmin:indmax,:);
div_vec = 1:(indmax-indmin+1);
 
n_cells = size(CDK2_normoxia,2);

figure
cell_age = [];
zz = 0;
clear CDK2_aligned_vec p21_aligned_vec cell_age
for k = 1:n_cells
    
divposns = [1 div_vec(cell_divisions_norm(:,k)==1) indmax-indmin 145];

CDK2_sc = [];
    for n = (length(divposns)-2)

        CDK2_smoothed_scdata_init = (smoothdata(CDK2_normoxia(divposns(n):divposns(n+1),k),'gaussian'));
        CDK2_smoothed_scdata = (smoothdata(CDK2_smooth(divposns(end):end,k),'gaussian'));
        P21_smoothed_scdata = (smoothdata(p21_smooth(divposns(end):end,k),'gaussian'));
        subplot(2,1,1)
        plotCDK2 = plot(0:1/3:(length(CDK2_smoothed_scdata)-1)/3, CDK2_smoothed_scdata);
        plotCDK2.Color(4) = 0.35;
        hold on
        subplot(2,1,2)
        plotP21 = plot(0:1/3:(length(P21_smoothed_scdata)-1)/3, P21_smoothed_scdata);
        plotP21.Color(4) = 0.35;
        hold on

        zz = zz + 1;
        CDK2_aligned_vec{zz} = (CDK2_normoxia(divposns(n):divposns(n+1),k));
        p21_aligned_vec{zz} = (p21_normoxia(divposns(n):divposns(n+1),k));
        cell_age = [cell_age; length(CDK2_aligned_vec{zz})];
        hold on

 
    end
end


%% Code for cell cycle classification and plotting both the Cdk2 trace that is used to classify cells as well as the whole CDK2 trace for that cell
for zz=1:3
    for k=1:n_cells
        if length(CDK2_aligned_vec{k})>0
            CDK2_aligned_trace =  CDK2_aligned_vec{k};
            cc_position(k) = cellcycleclassifier(CDK2_aligned_vec{k});

            if prctile(CDK2_aligned_trace,50)<0.38 && CDK2_aligned_trace(end)
            cc_position(k)=1;
            end

            if cc_position(k)==zz
            subplot(2,3,zz)
            ccplot = plot(((1:length(CDK2_aligned_vec{k}))-1)/3, CDK2_aligned_vec{k});
            hold on
            ccplot.Color(4) = 0.25;
            ylim([0 1])
            xlim([0 48])
            set(gca,'Xtick',[0 24 48])
            set(gca,'LineWidth',2)
            set(gca,'FontSize',18)
            set(gca,'Fontname','Arial')
            set(gca,'TickDir','out')
            if (zz == 1)
            ylabel('CDK2 activity')
            end
            pbaspect([1.1 1 1])
           
            
            subplot(2,3,zz+3)
            
            ccplot = plot((0:359)/3-48, CDK2_smooth(:,k)); 
            hold on
            ccplot.Color(4) = 0.25;
            ylim([0 1])
            xlim([-48 72])
            set(gca,'LineWidth',2)
            set(gca,'FontSize',18)
            set(gca,'Fontname','Arial')
            xlabel('Time (in h)')
            if (zz == 1)
            ylabel('CDK2 activity')
            end
            set(gca,'XTick',[-48 0 72])
            set(gca,'TickDir','out')
            pbaspect([1.1 1 1])
            end
        end
    end
end
        


%% Identifying sisters in cells right at hypoxia

% Code will not work properly after the cluster is discarded. Instead
% discard the cluster after sisters have been identified

sisters = [];

ndivs_percell = sum(cell_divisions(:,1:145)');
%We are going to take the progenitors at t=-48 (start of the experiment)
%and then find out which cells are sisters when they reach t = 0h at the
%start of hypoxic treatment


%First we consider cells that have divided only once. This is the easiest
%case because two cells with the same mother cell are sisters.
lineage_singlediv = lineagetree(ndivs_percell==1,1:145);

progenitors_singlediv = unique(lineage_singlediv(:,1));
singlediv_fam = zeros(length(progenitors_singlediv),2);
for k = 1:length(progenitors_singlediv)
[~,~,subfamily] = find(lineage_singlediv(lineage_singlediv(:,1)==progenitors_singlediv(k),end));
%if length(subfamily)==2
    singlediv_fam(k,:)=unique(subfamily);
%end
end


%Next we consider cells that have divided twice. In this case, we run a
%pairwise comparison of lineage overlap among all cells. The pairs of cells
%with the most lineage overlap are sisters. Others are cousins.
lineage_doublediv = lineagetree(ndivs_percell==2,1:145);
progenitors_doublediv = unique(lineage_doublediv(:,1));
doublediv_fam = zeros(length(progenitors_doublediv),4);

for k = 1:length(progenitors_doublediv)
    [~,~,subfamily_double] = find(lineage_doublediv(lineage_doublediv(:,1)==progenitors_doublediv(k),end));
    subfamily = unique(subfamily_double);
    if length(subfamily)<=4
    subfamily_padded = [subfamily', zeros(1,4-length(subfamily))];
    doublediv_fam(k,:) = subfamily_padded;
    clear subfamily subfamily_padded subfamily_double
    end
end

doublediv_fam(min(doublediv_fam')'==0,:) = [];
sisters = singlediv_fam;
cousins = [];
for k = 1:length(doublediv_fam)
    subfam = unique(doublediv_fam(k,:));
    subfam(subfam==0)=[];
    if length(subfam)==2
        sisters = [sisters; subfam];
    else
        if length(subfam)>2
        distpairs = zeros(length(subfam));
        for z = 1:length(subfam)-1
            for y = z+1:length(subfam)
                distpairs(z,y) = sum(lineagetree(subfam(z),1:145)==lineagetree(subfam(y),1:145));
            end
        end
        [m,I] = max(distpairs,[],'all','linear');
        sister1_pos = ceil(I/length(subfam));
        sister2_pos = mod(I,length(subfam));
        sisters = [sisters; subfam(sister1_pos), subfam(sister2_pos)];

        if length(subfam)>3
            distpairs(distpairs==m) = 0;
            [m2, I2] = max(distpairs,[],'all','linear');
            sister3_pos = ceil(I2/length(subfam));
            sister4_pos = mod(I2,length(subfam)); 
            sisters = [sisters; subfam(sister3_pos) subfam(sister4_pos)];

        end
        end
    end
end
            
        
%% Relative probability bar plots for sister cells identified previously. 
% The 2D histogram of cluster identities among sisters gives us
% probabilities P(Ti,Tj)) which is the probability of one sister cell
% belonging to cluster Ti while the other belongs to Tj. When i=j, they are
% in the same cluster (diagonal elements). All other cases are cells
% belonging to different clusters
T_new = T;
sis_hist = histcounts2(T_new(sisters(:,1)),T_new(sisters(:,2)));
sis_hist(1,:) = [];
sis_hist = sis_hist/(sum(sum(sis_hist)));
same_1 = sis_hist(1,1);
diff_1 = sum([sis_hist(1,:) sis_hist(:,1)'])-2*sis_hist(1,1);
same_2 = sis_hist(2,2);
diff_2 = sum([sis_hist(2,:) sis_hist(:,2)'])-2*sis_hist(2,2);
same_3 =  sis_hist(3,3);
diff_3 = sum([sis_hist(3,:) sis_hist(:,3)'])-2*sis_hist(3,3);
%Y = [same_1; same_2; same_3];
Y = [same_1/diff_1 same_2/diff_2 same_3/diff_3];
X = reordercats(categorical({'QQ','Q','P'}),{'QQ','Q','P'});
%X = categorical({'QQ','Q','P'});
bar(X,Y)
%%
% Shuffled cluster IDs for statistical testing
n_trials = 100000;
Y_shuffle = zeros(n_trials,3);
Y_same_shuffle = zeros(n_trials,1);
for k = 1:n_trials
 T_new_shuffled = T_new(randperm(length(T_new)));
sis_hist_shuffle = histcounts2(T_new_shuffled(sisters(:,1)),T_new_shuffled(sisters(:,2)),'Normalization','probability');

if size(sis_hist_shuffle,1)==4
    sis_hist_shuffle(1,:) = [];
end
if(size(sis_hist_shuffle,2)==4)
    sis_hist_shuffle(:,1) = [];
end
same_1 = sis_hist_shuffle(1,1);
diff_1 = sum([sis_hist_shuffle(1,:) sis_hist_shuffle(:,1)'])-2*sis_hist_shuffle(1,1);
same_2 = sis_hist_shuffle(2,2);
diff_2 = sum([sis_hist_shuffle(2,:) sis_hist_shuffle(:,2)'])-2*sis_hist_shuffle(2,2);
same_3 =  sis_hist_shuffle(3,3);
diff_3 = sum([sis_hist_shuffle(3,:) sis_hist_shuffle(:,3)'])-2*sis_hist_shuffle(3,3);
Y_shuffle(k,:) = [same_1/diff_1 same_2/diff_2 same_3/diff_3];
Y_same_shuffle(k) = same_1+same_2+same_3;
end

%% bar plot for cousins
T_new = T; 
T_new(T==3) = 1;
T_new(T==2) = 3;
T_new(T==1) = 2;
X = categorical({'Cluster QQ - Same','Cluster QQ - Different','Cluster Q - Same','Cluster Q - Different','Cluster P - Same', 'Cluster P - Different'});
sis_hist = histcounts2(T_new(cousins(:,1)),T_new(cousins(:,2)),'Normalization','probability');
same_1 = sis_hist(1,1);
diff_1 = sum([sis_hist(1,:) sis_hist(:,1)'])-2*sis_hist(1,1);
same_2 = sis_hist(2,2);
diff_2 = sum([sis_hist(2,:) sis_hist(:,2)'])-2*sis_hist(2,2);
same_3 =  sis_hist(3,3);
diff_3 = sum([sis_hist(3,:) sis_hist(:,3)'])-2*sis_hist(3,3);
Y = [same_1/diff_1 same_2/diff_2 same_3/diff_3];
X = reordercats(categorical({'QQ','Q','P'}),{'P','Q','QQ'});
bar(X,Y)


% Shuffled cluster IDs for statistical testing
n_trials = 100000;
Y_shuffle = zeros(n_trials,3);
for k = 1:n_trials
 T_new_shuffled = T_new(randperm(length(T_new)));
sis_hist_shuffle = histcounts2(T_new_shuffled(cousins(:,1)),T_new_shuffled(cousins(:,2)),(0:3),(0:3),'Normalization','probability');
same_1 = sis_hist_shuffle(1,1);
diff_1 = sum([sis_hist_shuffle(1,:) sis_hist_shuffle(:,1)'])-2*sis_hist_shuffle(1,1);
same_2 = sis_hist_shuffle(2,2);
diff_2 = sum([sis_hist_shuffle(2,:) sis_hist_shuffle(:,2)'])-2*sis_hist_shuffle(2,2);
same_3 =  sis_hist_shuffle(3,3);
diff_3 = sum([sis_hist_shuffle(3,:) sis_hist_shuffle(:,3)'])-2*sis_hist_shuffle(3,3);
Y_shuffle(k,:) = [same_1/diff_1 same_2/diff_2 same_3/diff_3];
end


%% plot the histograms
for k = 1:3
    for j = 1:3
        subplot(3,3,3*(k-1)+j)
        histogram(shufflehists(k,j,:),'Normalization','probability')
        hold on
        plot(nonshuffled_hist(k,j)*ones(1,2),[0 1])
    end
end

%% Function for cell cycle classification (Ref. Materials and Methods)

function z = cellcycleclassifier(input_trace)
    [~, max_ind] = max(input_trace);
    [~, min_ind] = min(input_trace);
    finalpt = length(input_trace);
    z=1;
    
    %G0/G1/S -> If x(end)<x(1) & minima is x(end), maxima is x(1)

    if(input_trace(end)<input_trace(1) && max_ind==1 && min_ind==finalpt)
        z = 1;
    end
    
    
    %G0/G1/S -> If minima is in between x(1) and x(end) and maxima is either x(1)
    %or x(end) and x(1)<x(end)
    if(input_trace(1)>input_trace(end) && min_ind~=1 && min_ind~=finalpt && (max_ind==1 || max_ind==finalpt))
        z = 1;
    end
    
    %S/G2 -> If minima is in between x(1) and x(end), maxima is either x(1) or x(end)
    %and x(end) > x(1)
    if(input_trace(end)>input_trace(1) && (max_ind==1 || max_ind==finalpt))
        z = 2;
    end
    
    %M -> If  maxima is in between x(1) and x(end)
    if(max_ind~=1 && max_ind~=finalpt)
        z = 3;
    end
    
    
    
end
```
