## Supplementary material for "Inertial effect of cell state velocity on the quiescence-proliferation fate decision in breast cancer": Code_Inertia_p21Cdk2.zip: detsoln_meanp21Cdk2vsCellCycleVelocity.m

```
clc
clear all


tic

%Define an empty vector for parameters
param = zeros(55,1);

%Define vectors for varying kSyE2f and kSyP21 by small amounts
%Define the number of times each parameter set will be simulated with
%slightly randomized initial conditions
mod_vec_2 = 0:0.05:1;
mod_vec = -0.1:0.05:0.2;
ntrials = 10;

numdatapts = length(mod_vec_2)*length(mod_vec)*ntrials;

%Define empty matrices for storing s-phase entry times and mean p21, Cdk2
%values
t_sphase_vec = zeros(1,numdatapts);
t_sphase_end_vec = zeros(1,numdatapts);

aCdk2_vec = zeros(1,numdatapts);
tP21_vec = zeros(1,numdatapts);
aCdk2_preS_vec = zeros(1,numdatapts);
tP21_preS_vec = zeros(1,numdatapts);
aCdk2_G2_vec = zeros(1,numdatapts);
tP21_G2_vec = zeros(1,numdatapts);
loopvar = 0;
for mm = 1:length(mod_vec_2)
for kk = 1:length(mod_vec)

%Each parameter set is simulated multiple times to account for
%stochasticity in initial p21-Cdk2 values
for nn = 1:ntrials
    loopvar = loopvar+1;

%Define parameters and then vary kSyE2f and kSyP21
kSyE2f    = 0.03*10^mod_vec(kk);  %AU/min  constitutive E2F synthesis
kSyE2fE2f = 0.04    ;   %AU/min  E2F-dependent E2F synthesis
jSyE2f    = 0.2      ;  %AU  Michealis-Menten constant for E2F synthesis
kAsRbE2f  = 5          ; %1/(AUmin)  association of Rb and E2F
kDsRbE2f  = 0.005     ;%1/min  dissociation of Rb:E2F complexes
kDeE2f    = 0.05    ; %1/min  constiutive E2F degradation

kPhRbCd   = 0.2     ;%1/(AUmin)  CycD:CDK4/6-mediated Rb phosphorylation
kPhRbCe   = 0.3      ; %1/(AUmin)  CycE:CDK2-mediated Rb phosphorylation
kPhRbCa   = 0.3      ;%1/(AUmin)  CycA:Cdk2-mediated Rb phosphorylation
kDpRb     = 0.05    ;   %1/min  dephosphorylation of Rb

kSyE1   = 0.005       ;  %1/min  constitutive Emi1 synthesis
kDeE1C1 = 0.005    ; %1/min  Cdh1-mediated Emi1 degradation
kDeE1   = 0.0005       ;%1/min  constitutive Emi1 degradation

kPhC1   = 0            ;%1/min  constitutive Cdh1 phosphorylation
kPhC1Ce = 0.01     ;%1/(AUmin)  CycE-mediated Cdh1 phosphorylation
kPhC1Ca = 1       ;%1/(AUmin)  CycA-mediated Cdh1 phosphorylation
kDpC1   = 0.05        ;%1/min  Cdh1 dephosphorylation
kAsE1C1 = 10         ;%1/(AUmin)  association of Emi1 and Cdh1
kDsE1C1 = 0.01       ;%1/min  dissociation of Emi1:Cdh1 complexes

%kSyP21 variation
kSyP21    = 0.002*10^mod_vec_2(mm)   ;   %AU/min  constitutive p21 synthesis
kSyP21P53 = 0.008     ;  %1/min  p53-dependent synthesis of p21
kDeP21    = 0.0025     ; %1/min  constitutive p21 degradation
kDeP21Cy  = 0.007       ;%1/(AUmin)  cyclin:CDK2-mediated p21 degradation (Skp2-dependent)
kDeP21aRc = 1           ;%1/(AUmin)  aRC-mediated p21 degradation (Cdt2-dependent)

kSyCe     = 0.01       ; %1/min  constitutive CycE synthesis
kSyCa     = 0.02        ;%1/min  constitutive CycA synthesis
kAsCyP21  = 1          ; %1/(AUmin)  association of cyclin:CDK2 and p21
kDsCyP21  = 0.05        ;%1/min  dissociation of cyclin:CDK2:p21 complexes
kDeCe     = 0.004       ;%1/min  constitutive CycE degradation
kDeCa     = 0.01        ;%1/min  constitutive CycA degradation
kDeCeCa   = 0.015       ;%1/(AUmin)  CycA:CDK2-mediated CycE degradation (Skp2-dependent)
kDeCaC1   = 2            ;%1/(AUmin)  Cdh1-mediated CycA degradation

kImPc     = 0.003       ; %AU/min  PCNA import into the nucleus
kExPc     = 0.006       ; %1/min   PCNA export from the nucleus

kPhRc     = 0.1          ;%1/min  cyclin:CDK2-mediated phosphorylation/priming of RCs
kDpRc     = 0.05         ;%1/min  dephosphorylation of pRCs
jCy       = 1.8          ;%AU  CDK2 threshold for the phosphorylation/priming of RCs
n         = 6            ;%-  hill coefficient for the phosphorylation/priming of RCs
kAsRcPc   = 0.01         ;%1/(AUmin)  association of phosphorylated/primed RCs and PCNA
kDsRcPc   = 0.001        ;%1/min  dissociation of pRC:PCNA complexes
kAsPcP21  = 100           ;%1/(AUmin)  association of PCNA and p21
kDsPcP21  = 0.01         ;%1/min  dissociation of PCNA:p21 complexes
kSyDna    = 0.0093       ;%1/min  DNA synthesis by aRCs

kSyP53    = 0.05         ;%AU/min  constitutive p53 synthesis
kDeP53    = 0.05         ;%AU/min  DNA damage-dependent degradation of p53
jP53      = 0.01         ;%AU  inhibition constant of p53 degradation by DNA damage


kReDam    = 0.001        ;%1/min  p53-independent DNA damage repair
kReDamP53 = 0.005        ;%1/min  p53-dependent DNA damage repair
jDam      = 0.5         ;%AU  DNA damage threshold for repair


Cd   = 0.65; 	%AU  relative CycD:Cdk4/6 level
tRb  = 5        ;     	%AU  total level of Rb
tC1  = 1         ;       %AU  total APC/C^Cdh1 level
Skp2 = 1          ;      %-  relative SCF^Skp2 level
Cdt2 = 1           ;     %-  relative CRL4^Cdt2 level


rDsRc     = 1; %Disassembly of DNA complexes at end of S phase


y_init = zeros(1,20);

%Initial conditions from Heldt et al., varied using a normal distribution
%with 20% variance.
y_init(1,5) = max(0,0.6*(1+randn*0.2));%tP21
y_init(1,6) = max(0,0.5*(1+randn*0.2)); %tCe

y_init(1,7) = max(0,1.2*(1+randn*0.2)); %tCa

y_init(1,12) = 0.5;%aPcna
y_init(1,14) = 1; %Rc


%Parameters
param(1)=kSyE2f;
param(2)=kSyE2fE2f;
param(3)=jSyE2f;
param(4)=kAsRbE2f;
param(5)=kDsRbE2f;
param(6)=kDeE2f;
param(7)=kPhRbCd;
param(8)=kPhRbCe;
param(9)=kPhRbCa;
param(10)=kDpRb;
param(11)=kSyE1;
param(12)=kDeE1C1 ;
param(13)=kDeE1 ;
param(14)=kPhC1 ;
param(15)=kPhC1Ce ;
param(16)=kPhC1Ca ;
param(17)=kDpC1;
param(18)=kAsE1C1;
param(19)=kDsE1C1;
param(20)=kSyP21 ;
param(21)=kSyP21P53 ;
param(22)=kDeP21 ;
param(23)=kDeP21Cy;
param(24)=kDeP21aRc;
param(25)=kSyCe ;
param(26)=kSyCa ;
param(27)=kAsCyP21 ;
param(28)=kDsCyP21;
param(29)=kDeCe ;
param(30)=kDeCa;
param(31)=kDeCeCa ;
param(32)=kDeCaC1;
param(33)=kImPc;
param(34)=kExPc ;
param(35)=kPhRc;
param(36)=kDpRc ;
param(37)=jCy;
param(38)=n ;
param(39)=kAsRcPc;
param(40)=kDsRcPc;
param(41)=kAsPcP21;
param(42)=kDsPcP21;
param(43)=kSyDna;
param(44)=kSyP53;
param(45)=kDeP53;
param(46)=jP53;
param(47)=kReDam;
param(48)=kReDamP53;
param(49)=jDam ;

%Other global variables
param(50)=Cd*(1+randn*0.2);
param(51)=tRb;
param(52)=tC1;
param(53)=Skp2;
param(54)=Cdt2;

param(55)=rDsRc;


tspan_1 = (0:0.1:2000);

[t_1, y_1] = ode23s(@(t,y) detsys_meanp21Cdk2(t,y,param), tspan_1, y_init);

if(y_1(end,18)>0.9)
    %Find when S-phase begins
[~, t_sphase_ind] = min(abs(y_1(:,18) - 0.1));

t_sphase = interp1([y_1(t_sphase_ind-1,18) y_1(t_sphase_ind,18) y_1(t_sphase_ind+1,18)],[t_1(t_sphase_ind-1) t_1(t_sphase_ind) t_1(t_sphase_ind+1)],0.1);

[~, t_sphase_end_ind] = min(abs(y_1(t_sphase_ind:end,16)-0.1));
t_sphase_end_ind = t_sphase_end_ind + t_sphase_ind - 1;

if t_sphase_end_ind < length(t_1)
t_sphase_end = interp1([y_1(t_sphase_end_ind-1,16) y_1(t_sphase_end_ind,16) y_1(t_sphase_end_ind+1,16)],[t_1(t_sphase_end_ind-1) t_1(t_sphase_end_ind) t_1(t_sphase_end_ind+1)],0.1,'linear','extrap');
else
    t_sphase_end = t_1(end);
end

t_sphase_vec(loopvar) = t_sphase;

t_sphase_end_vec(loopvar) = t_sphase_end;

y_init_2 = y_1(t_sphase_end_ind,:);
%Simulate for an extra 4 hours to be representative of G2 that is present
%in experimental data
tspan_2 = ((t_sphase_end):0.1:(t_sphase_end+240));

[t_2, y_2] = ode23s(@(t,y) detsys_meanp21Cdk2(t,y,param), tspan_2, y_init_2);

aCdk2_G2_vec(loopvar) = mean(y_1(1:t_sphase_ind,6) + y_1(1:t_sphase_ind,7) - y_1(1:t_sphase_ind,8) - y_1(1:t_sphase_ind,9));
tP21_G2_vec(loopvar) = mean(y_1(1:t_sphase_ind,5));

else

aCdk2_G2_vec(loopvar) = mean(y_1(:,6) + y_1(:,7) - y_1(:,8) - y_1(:,9));
tP21_G2_vec(loopvar) = mean(y_1(:,5));

   t_sphase_vec(loopvar) = Inf; 
   t_sphase_end_vec(loopvar) = Inf;
end
aCdk2_vec(loopvar) = mean(y_1(:,6) + y_1(:,7) - y_1(:,8) - y_1(:,9));
tP21_vec(loopvar) = mean(y_1(:,5));
end


end
end


%% Plotting meanp21 and mean Cdk2 against cell cycle velocity

%Considering only proliferative cells
tP21_prolif = tP21_G2_vec(t_sphase_vec~=Inf);
aCdk2_prolif = aCdk2_G2_vec(t_sphase_vec~=Inf);

t_sphase_end_prolif = t_sphase_end_vec(t_sphase_vec~=Inf);

%Since this is a scatter plot and it is difficult to simply get a mean, we
%bin the cell cycle velocities instead and then calculate a mean and
%percentile range for each bin
nbins = 10;
[~, edges, binID] = histcounts(1./t_sphase_end_prolif,nbins);

for k = 1:nbins
    if(sum(binID==k))>5
    tP21_mean(k) = mean(tP21_prolif(binID==k));
    tP21_90(k) = prctile(tP21_prolif(binID==k),90);
    tP21_10(k) = prctile(tP21_prolif(binID==k),10);
    
    aCdk2_mean(k) = mean(aCdk2_prolif(binID==k));
    aCdk2_90(k) = prctile(aCdk2_prolif(binID==k),90);
    aCdk2_10(k) = prctile(aCdk2_prolif(binID==k),10);
    else
    tP21_mean(k) = NaN;
    tP21_90(k) = NaN;
    tP21_10(k) = NaN; 
    
    aCdk2_mean(k) = NaN;
    aCdk2_90(k) = NaN;
    aCdk2_10(k) = NaN;
    end
end


figure
scatter(1./t_sphase_end_vec, tP21_G2_vec)
hold on
plot((edges(1:end-1)+edges(2:end))/2, tP21_mean,'-o')
plot((edges(1:end-1)+edges(2:end))/2, tP21_90)
plot((edges(1:end-1)+edges(2:end))/2, tP21_10)
set(gca,'Yscale','log')
xlim([-0.5 4.5]*10^-3)
ylim(10.^[-1 1])
set(gca,'TickDir','out')
set(gca,'LineWidth',2)
set(gca,'FontSize',24)
set(gca,'Fontname','arial')
box on


figure
scatter(1./t_sphase_end_vec, aCdk2_G2_vec./(aCdk2_G2_vec + 1))
hold on
plot((edges(1:end-1)+edges(2:end))/2, aCdk2_mean./(aCdk2_mean+1),'-o')
plot((edges(1:end-1)+edges(2:end))/2, aCdk2_90./(aCdk2_90+1))
plot((edges(1:end-1)+edges(2:end))/2, aCdk2_10./(aCdk2_10+1))
xlim([-0.5 4.5]*10^-3)
ylim([0 0.8])
ytickformat('%.1f')
set(gca,'TickDir','out')
set(gca,'LineWidth',2)
set(gca,'FontSize',24)
set(gca,'Fontname','arial')
box on
```
