## Supplementary material for "Inertial effect of cell state velocity on the quiescence-proliferation fate decision in breast cancer": Code_Inertia_p21Cdk2.zip: detsys_meanp21Cdk2.m

```
function dydt = detsys_meanp21Cdk2(t,y, param)


uRb = y(1);
tE2f = y(2);
E2f = y(3);
tE1 = y(4);
tP21 = y(5);
tCe = y(6);
tCa = y(7);

CeP21 = y(8);
CaP21 = y(9);

C1 = y(10);
E1C1 = y(11);

aPcna = y(12);
iPcna = y(13);

Rc = y(14);
pRc = y(15);
aRc = y(16);
iRc = y(17);

Dna = y(18);

P53 = y(19);
Dam = 0;


%Parameters
kSyE2f = param(1);
kSyE2fE2f = param(2);
jSyE2f = param(3);
kAsRbE2f = param(4);
kDsRbE2f = param(5);
kDeE2f = param(6);


kPhRbCd = param(7);
kPhRbCe = param(8);
kPhRbCa = param(9);
kDpRb = param(10);     

kSyE1 = param(11);  
kDeE1C1  = param(12);
kDeE1  = param(13); 

kPhC1  = param(14);
kPhC1Ce  = param(15);
kPhC1Ca  = param(16);
kDpC1 = param(17);
kAsE1C1 = param(18);
kDsE1C1 = param(19);

kSyP21  = param(20);
kSyP21P53  = param(21);
kDeP21  = param(22);   
kDeP21Cy = param(23);
kDeP21aRc = param(24);

kSyCe  = param(25);
kSyCa  = param(26);
kAsCyP21  = param(27);
kDsCyP21 = param(28);
kDeCe  = param(29);
kDeCa   = param(30);
kDeCeCa  = param(31);
kDeCaC1   = param(32);

kImPc   = param(33);  
kExPc    = param(34);

kPhRc = param(35);
kDpRc  = param(36);
jCy   = param(37);
n  = param(38);
kAsRcPc = param(39);
kDsRcPc = param(40);
kAsPcP21 = param(41);
kDsPcP21 = param(42);
kSyDna  = param(43);

kSyP53  = param(44);
kDeP53  = param(45);
jP53   = param(46);  

kReDam   = param(47);
kReDamP53  = param(48);
jDam  = param(49);    

%Other global variables
Cd = param(50);
tRb = param(51);
tC1 = param(52);
Skp2 = param(53);
Cdt2 = param(54);

rDsRc = param(55);


Rb    = uRb  - tE2f  + E2f;      %AU  free, unphoshporylated Rb
pRb   = tRb  - uRb         ;     %AU  free, phosphorylated Rb
RbE2f = tE2f - E2f          ;    %AU  Rb:E2F complexes

E1  = tE1  - E1C1            ;               %AU  free Emi1
pC1 = tC1  - C1 - E1C1        ;              %AU  free, phosphorylated Cdh1
Ce  = tCe  - CeP21             ;             %AU  free, active CycE
Ca  = tCa  - CaP21              ;            %AU  free, active CycA
P21 = tP21 - CeP21 - CaP21 - iPcna - iRc;    %AU  free p21

rSyE2f = kSyE2f + kSyE2fE2f*E2f / (jSyE2f + E2f);               %AU/min E2F synthesis
rPhRb  = kPhRbCd*Cd + kPhRbCe*Ce + kPhRbCa*Ca    ;               %1/min  phosphorylation of Rb
rPhRc  = kPhRc  *(Ce+Ca)^n / (jCy^n+(Ce+Ca)^n)    ;              %1/min  phosphorylation/priming of replication complexes
rPhC1  = kPhC1 + kPhC1Ce*Ce + kPhC1Ca*Ca           ;             %1/min  Cdh1 phosphorylation

rDeP21 = kDeP21 + kDeP21Cy*Skp2*(Ce+Ca) + kDeP21aRc*Cdt2*aRc; 	%1/min  p21 degradation
rDeCe  = kDeCe  + kDeCeCa*Ca                                 ;	%1/min  CycE degradation 
rDeCa  = kDeCa  + kDeCaC1*C1                                ;	%1/min  CycA degradation

rDeP53 = kDeP53 / (jP53 + Dam) ;                                 % p53 degradation
rReDam = kReDam + kReDamP53*P53 / (jDam + Dam)     ;             % DNA damage repair

dydt(1)= -rPhRb*uRb+kDpRb*pRb;
dydt(2) = (rSyE2f - kDeE2f*tE2f);
dydt(3)  = rSyE2f - kDeE2f*E2f - kAsRbE2f*Rb*E2f + (kDsRbE2f+rPhRb)*RbE2f;

dydt(4)  = (kSyE1 *E2f - kDeE1C1*E1C1  - kDeE1*E1);
dydt(5) = kSyP21     + kSyP21P53*P53 - rDeP21*tP21;
dydt(6)  = kSyCe *E2f                 - rDeCe*tCe;
dydt(7)  = kSyCa *E2f                 - rDeCa*tCa;

dydt(8) = kAsCyP21*P21*Ce - (kDsCyP21+rDeCe+rDeP21)*CeP21;
dydt(9) = kAsCyP21*P21*Ca - (kDsCyP21+rDeCa+rDeP21)*CaP21;

dydt(10)  = - rPhC1*C1 + kDpC1*pC1 - kAsE1C1*E1*C1 + (kDsE1C1+kDeE1C1)*E1C1;
dydt(11) = - rPhC1*E1C1           + kAsE1C1*E1*C1 - (kDsE1C1+kDeE1C1)*E1C1;

dydt(12) = kImPc - kAsPcP21*P21*aPcna + (kDsPcP21+rDeP21)*iPcna - (kExPc+kAsRcPc*pRc)*aPcna + (kDsRcPc+rDsRc)*aRc;
dydt(13) =       + kAsPcP21*P21*aPcna - (kDsPcP21+rDeP21)*iPcna - (kExPc+kAsRcPc*pRc)*iPcna + (kDsRcPc+rDsRc)*iRc;

if Dna<1
dydt(14)  = - rPhRc*Rc + kDpRc*pRc;
dydt(15) =   rPhRc*Rc - kDpRc*pRc                     - kAsRcPc*(aPcna+iPcna)*pRc + kDsRcPc*(aRc+iRc);
dydt(16) = - kAsPcP21*P21*aRc + (kDsPcP21+rDeP21)*iRc + kAsRcPc* aPcna       *pRc - kDsRcPc*aRc;
dydt(17) =   kAsPcP21*P21*aRc - (kDsPcP21+rDeP21)*iRc + kAsRcPc*       iPcna *pRc - kDsRcPc*iRc;
else
dydt(14)  = - rPhRc*Rc + kDpRc*pRc                                                                     - rDsRc*Rc;
dydt(15) =   rPhRc*Rc - kDpRc*pRc                     - kAsRcPc*(aPcna+iPcna)*pRc + kDsRcPc*(aRc+iRc) - rDsRc*pRc;
dydt(16) = - kAsPcP21*P21*aRc + (kDsPcP21+rDeP21)*iRc + kAsRcPc* aPcna       *pRc - kDsRcPc*aRc       - rDsRc*aRc;
dydt(17) =   kAsPcP21*P21*aRc - (kDsPcP21+rDeP21)*iRc + kAsRcPc*       iPcna *pRc - kDsRcPc*iRc       - rDsRc*iRc;
end

if Dna<1
dydt(18) = kSyDna*aRc;
else
dydt(18) = 0;
end


dydt(19) = kSyP53 - rDeP53*P53;
%Activate below in case of DNA damage
%dydt(20) = kGeDam + kGeDamArc*aRc - rReDam*Dam;
%dydt(20) = 0.001 + 0.012*aRc - (kReDam + kReDamP53*(P53)/(jDam+Dam))*Dam;
dydt(20) = 0;


dydt = dydt';

end
```
