## Supplementary material for "Inertial effect of cell state velocity on the quiescence-proliferation fate decision in breast cancer": Code_Inertia_p21Cdk2.zip: File descriptions

Description of supplementary computational files


%%% Image analysis and single cell tracking %%%

p21Cdk2Measurements_Hypoxia.mat - contains the raw trace values from p53Cinema analysis

AnalysisScript_p21Cdk2Traces.m - script file used for all calculations done in the paper (Smoothing, clustering, cell cycle analysis, sister cell analysis)


%%% Energy landscape generation %%%

reactionnetwork_stoch.jl - Julia file for the stochastic reaction network from Heldt et al. (PNAS 2018)

p21CDK2landscape.jl - Julia file for generating trajectories of Heldt et al. (PNAS 2018) model

tall_landscapegen.m - MATLAB file for visualization of the energy landscape in terms of total p21 and active Cdk2


%%% Cell cycle velocity simulations %%%

SPhaseEntry_SensitivityAnalysis.m - Sensitivity analysis for S phase entry time

detsys_Sphasetiming.m - Function file containing the Heldt et al. model used in SPhaseEntry_SensitivityAnalysis.m


detsoln_meanp21Cdk2vsCellCycleVelocity.m - Calculation and plotting of mean total p21 and mean active Cdk2 as a function of cell cycle velocity

detsys_meanp21Cdk2.m - Function file containing Heldt et al. model used in detsoln_meanp21Cdk2vsCellCycleVelocity.m 
