## Supplementary material for "Inertial effect of cell state velocity on the quiescence-proliferation fate decision in breast cancer": Code_Inertia_p21Cdk2.zip: p21CDK2landscape.jl

```
#This code is for generation of an energy landscape by randomizing initial conditions of p21 and Cdk2 associated species

#Section 1 - Initializing parallelization. Beyond this, the @everywhere macro is to perform instructions on ALL the workers
using Distributed

if nworkers()==1
addprocs(15)
end

#Section 1 - end


#Section 2 - Initializing packages for simulation
@time begin
@everywhere using DifferentialEquations, Plots, Distributed, StaticArrays, DataFrames, CSV, ModelingToolkit, Catalyst
end

#Section 2 - end

#Section 3 - Setting up the simulations

#Loading the p21-Cdk2 reaction network
@everywhere include("reactionnetwork_stoch.jl");

@everywhere begin
    
#Initializing all parameters
Cd	=	650;
Skp2	=	1	;
Cdt2	=	1	;
kSyE2f	=	30	;
kSyE2fE2f	=	40	;
jSyE2f	=	200	;
kAsRbE2f	=	0.005	;
kDsRbE2f	=	0.005	;
kDeE2f	=	0.05	;
kPhRbCd	=	0.0002	;
kPhRbCe	=	0.0003	;
kPhRbCa	=	0.0003	;
kDpRb	=	0.05	;
kSyE1	=	0.005	;
kDeE1C1	=	0.005	;
kDeE1	=	0.0005	;
kPhC1	=	0	;
kPhC1Ce	=	1.00E-05	;
kPhC1Ca	=	0.001	;
kDpC1	=	0.05	;
kAsE1C1	=	0.01	;
kDsE1C1	=	0.01	;
kSyP21	=	2	;
kSyP21P53	=	0.008	;
kDeP21	=	0.0025	;
kDeP21Cy	=	7.00E-06	;
kDeP21aRc	=	0.001	;
kSyCe	=	0.01	;
kSyCa	=	0.02	;
kAsCyP21	=	0.001	;
kDsCyP21	=	0.05	;
kDeCe	=	0.004	;
kDeCa	=	0.01	;
kDeCeCa	=	1.50E-05	;
kDeCaC1	=	0.002	;
kImPc	=	3	;
kExPc	=	0.006	;
kPhRc	=	0.1	;
kDpRc	=	0.05	;
jCy	=	1800	;
n	=	6	;
kAsRcPc	=	1.00E-05	;
kDsRcPc	=	0.001	;
kAsPcP21	=	0.1	;
kDsPcP21	=	0.01	;
kDeP53	=	5.0	;
    kSyP53 = 50.0
#Initial conditions, recast as a vector, randomized for p21 and Cdk2
        u0 = [0, 0, 0, Int64(round(rand()*1800)), Int64(round(rand()*1500)), 0, Int64(round(rand()*1000)), 0, 0, 1000, 0, 0, 0, 5000, 0, 0, 0, 0, 0, 0];
#Parameter vection
    p = [Cd, Skp2, Cdt2, kSyE2f, kSyE2fE2f, jSyE2f, kAsRbE2f, kDsRbE2f, kDeE2f, kPhRbCd, kPhRbCe, kPhRbCa, kDpRb, kSyE1, kDeE1C1, kDeE1, kPhC1, kPhC1Ce, kPhC1Ca, kDpC1, kAsE1C1, kDsE1C1, kSyP21, kSyP21P53, kDeP21, kDeP21Cy, kDeP21aRc, kSyCe, kSyCa, kAsCyP21, kDsCyP21, kDeCe, kDeCa, kDeCeCa, kDeCaC1, kImPc, kExPc, kPhRc, kDpRc, jCy, n, kAsRcPc, kDsRcPc, kAsPcP21, kDsPcP21, kSyP53, kDeP53];# kSyP53, kDeP53, jP53];
#Time span of the simulation
    tspan = (0.0, 10000.0);
#Generate a discrete problem where you are working in molecules instead of concentrations
    dprob = DiscreteProblem(rn,u0,tspan,p);
#Convert of a jump system so you can use with Gillespie
    jsys = convert(JumpSystem,rn);
#Recast as a problem so you can solve discrete problem as a jump system. 
#This can be solved directly as a discrete problem, but recasting it as a jump problem lets us set save_positions to false
#allowing us to then reduce memory usage
    jprob = JumpProblem(jsys, dprob, SortingDirect(); save_positions=(false,false));
end


#Generate a probability function that randomizes inputs so that each run starts with random inputs
@everywhere begin
function prob_func(jprob,i,repeat)
        pRb::Int64 = 0;
        
        C1::Int64 = 1000;
        pC1::Int64 = 0;
        E1C1::Int64 = 0;
        E1::Int64 = 0;
        #E1::Int64 = 0;

        tE2f::Int64 = 0;
        RbE2f::Int64 =0;
        Rb::Int64 =  5000;
        E2f::Int64 = 0;

        tCdk2::Int64 = Int64(round(3000*rand()));
        tCa::Int64  = Int64(round(tCdk2*rand()));
        tCe::Int64  = (tCdk2 - tCa);

        tP21::Int64  = Int64(round(1000*rand()));

        CaP21::Int64 = Int64(round(rand()*min(tCa,tP21)));
        CeP21::Int64 = Int64(round(rand()*min(tCe,(tP21 - CaP21))));

        Ca::Int64 = tCa - CaP21;
        Ce::Int64 = tCe - CeP21;
        
        Rc::Int64 = 1000;
        pRc::Int64 =0;
        iRc::Int64 = 0;
        aRc::Int64 = 0;

        aPcna::Int64 = 500;
        iPcna::Int64 = 0;
                
        P21::Int64 = tP21 - CeP21 - CaP21 - iPcna - iRc;
        
        p53::Int64 = Int64(round(rand()*3000));
        

@. jprob.prob.u0 = Int64.(round.([aPcna, pRc, aRc, Ca, P21, CaP21, Ce, CeP21, E1, C1, E1C1, iPcna, iRc, Rb, E2f, RbE2f, pC1, pRb, Rc, p53]));

    jprob
end
# Ensemble problem for parallelization of the simulations to run in Julia
     eprob = EnsembleProblem(jprob; prob_func = prob_func);#, output_func = output_func);#, #prob_func=prob_func);
end

#Output file for the trajectories
outputfile = string("gillespie_paths_Cd.csv");

#Section 3 - end


#Section 4 - Running the simulation

#We set up a for loop that runs 200 trajectories each time with 16 cores, saving at 0.5
#After running and saving, we flush stdout and run garbage collection
#Running everything together without batching, garbage collection, and storage caused out-of-memory issues even with a lot of RAM (~128 GB)
for k = 1:25
ntrajs = 200;
sol = solve(eprob, SSAStepper(), EnsembleDistributed(); trajectories = ntrajs, batch_size = 16, saveat=0.5);
flush(stdout)
for zz = 1:ntrajs
CSV.write(outputfile,sol[zz],append=true);
    end
println("Done running $(ntrajs*k)")
        flush(stdout)
       GC.gc()
end
```
