## Supplementary material for "Inertial effect of cell state velocity on the quiescence-proliferation fate decision in breast cancer": Code_Inertia_p21Cdk2.zip: reactionnetwork_stoch.jl

```
#p21Cdk2 reaction network for stochastic simulations


rn = @reaction_network begin
	kAsRcPc, aPcna + pRc --> aRc #Association of active PCNA and replication complexes
	kAsCyP21, Ca + P21 --> CaP21 #Association of CycA:Cdk2 and p21
	kAsCyP21, Ce + P21 --> CeP21 #Association of CycE:Cdk2 and p21
	kAsE1C1, E1 + C1 --> E1C1 #Association of Emi1:APC/C^Cdh1 complexes
	kAsRcPc, iPcna + pRc --> iRc #Association of inactive PCNA and replication complexes
	kAsPcP21, aRc + P21 --> iRc #Association of p21 and replication complexes
	kAsPcP21, aPcna + P21 --> iPcna #Association of PCNA and p21
	kAsRbE2f, Rb + E2f --> RbE2f #Association of Rb and E2F
	#Reaction with degradation of probe neglected
	(kDeCa + kDeCaC1*C1), Ca --> 0 #Degradation of CycA in CycA:Cdk2 complexes
	(kDeCa + kDeCaC1*C1), CaP21 --> P21 #Degradation of CycA in CycA:Cdk2:p21 complexes
	(kDeCe + kDeCeCa*Ca), Ce --> 0 #Degradation of CycE in CycE:Cdk2 complexes
	(kDeCe + kDeCeCa*Ca), CeP21 --> P21 #Degradation of CycE in CycE:Cdk2:p21 complexes
	kDeE2f, E2f --> 0 #Degradation of E2F
	kDeE2f, RbE2f --> Rb #Degradation of E2F in Rb:E2F complexes
	kDeE1, E1 --> 0 #Degradation of Emi1
	kDeE1C1, E1C1 --> C1 #Degradation of Emi1 in Emi1:APC/C^Cdh1 complexes
	(kDeP21 + kDeP21Cy*Skp2*(Ce+Ca) + kDeP21aRc*Cdt2*aRc), P21 --> 0 #Degradation of free p21
	(kDeP21 + kDeP21Cy*Skp2*(Ce+Ca) + kDeP21aRc*Cdt2*aRc), CaP21 --> Ca #Degradation of p21 in CycA:Cdk2:p21 complexes
	(kDeP21 + kDeP21Cy*Skp2*(Ce+Ca) + kDeP21aRc*Cdt2*aRc), CeP21 --> Ce #Degradation of p21 in CycE:Cdk2:p21 complexes
	(kDeP21 + kDeP21Cy*Skp2*(Ce+Ca) + kDeP21aRc*Cdt2*aRc), iRc --> aRc #Degradation of p21 in inactive replication complexes
	(kDeP21 + kDeP21Cy*Skp2*(Ce+Ca) + kDeP21aRc*Cdt2*aRc), iPcna --> aPcna #Degradation of p21 in PCNA:p21 complexes
	kDpC1, pC1 --> C1 #Dephosphorylation of APC/C^Cdh1
	kDpRb, pRb --> Rb #Dephosphorylation of Rb
	kDpRc, pRc --> Rc #Dephosphorylation of replication complexes
	#Neglecting disassembly of replication complexes
	kDsRcPc, aRc --> aPcna + pRc #Dissociation of active PCNA and replication complexes
	kDsCyP21, CaP21 --> Ca + P21 #Dissociation of CycA:Cdk2 and p21
	kDsCyP21, CeP21 --> Ce + P21 #Dissociation of CycE:Cdk2 and p21
	kDsE1C1, E1C1 --> E1 + C1 #Dissociation of Emi1:APC/C^Cdh1 complexes
	kDsRcPc, iRc --> iPcna + pRc #Dissociation of inactive PCNA and replication complexes
	kDsPcP21, iRc --> aRc + P21 #Dissociation of p21 and replication complexes
	kDsPcP21, iPcna --> aPcna + P21 #Dissociation of PCNA and p21
	kDsRbE2f, RbE2f --> Rb + E2f #Dissociation of Rb and E2F
	#Neglecting induction of DNA damage
	kExPc, aPcna --> 0 #Nuclear export of active PCNA
	kExPc, iPcna --> P21 #Nuclear export of inactive PCNA
	kImPc, 0 --> aPcna #Nuclear import of active PCNA
	(kPhC1 + kPhC1Ce*Ce + kPhC1Ca*Ca), E1C1 --> E1 + pC1 #Phosphorylation of APC/C^Cdh1 in Emi1:APC/C^Cdh1 complexes
	(kPhC1 + kPhC1Ce*Ce + kPhC1Ca*Ca), C1 --> pC1 #Phosphorylation of free APC/C^Cdh1
	(kPhRbCd*Cd + kPhRbCe*Ce + kPhRbCa*Ca), Rb --> pRb #Phosphorylation of Rb
	(kPhRbCd*Cd + kPhRbCe*Ce + kPhRbCa*Ca), RbE2f --> pRb + E2f #Phosphorylation Rb in Rb:E2F complexes
	((kPhRc*(Ce+Ca)^n)/(jCy^n + (Ce + Ca)^n)), Rc --> pRc #Phosphorylation/priming of replication complexes
	#Neglecting DNA damage repair
	#Neglecting probe synthesis
	kSyCa, E2f --> E2f + Ca #Synthesis of CycA
	kSyCe, E2f --> E2f + Ce #Synthesis of CycE
	#Neglecting DNA synthesis
	(kSyE2f + (kSyE2fE2f*E2f/(jSyE2f + E2f))), 0 --> E2f #Synthesis of E2F
	kSyE1, E2f --> E2f + E1 #Synthesis of Emi1
	(kSyP21 + kSyP21P53*P53), 0 --> P21 #Synthesis of P21
	kSyP53, 0 --> P53 #Synthesis of P53
    kDeP53, P53 --> 0 #Degradation of P53
end Cd Skp2 Cdt2 kSyE2f kSyE2fE2f jSyE2f kAsRbE2f kDsRbE2f kDeE2f kPhRbCd kPhRbCe kPhRbCa kDpRb kSyE1 kDeE1C1 kDeE1 kPhC1 kPhC1Ce kPhC1Ca kDpC1 kAsE1C1 kDsE1C1 kSyP21 kSyP21P53 kDeP21 kDeP21Cy kDeP21aRc kSyCe kSyCa kAsCyP21 kDsCyP21 kDeCe kDeCa kDeCeCa kDeCaC1 kImPc kExPc kPhRc kDpRc jCy n kAsRcPc kDsRcPc kAsPcP21 kDsPcP21 kSyP53 kDeP53
```
