## Supplementary material for "Inertial effect of cell state velocity on the quiescence-proliferation fate decision in breast cancer": Code_Inertia_p21Cdk2.zip: SPhaseEntry_SensitivityAnalysis.m

```
tic
clear all

%% Latin hypercube based simulations for S-phase timing
 
num_scens = 1000; %number of sample values (or scenarios)
p = 55; %number of variables
%returns an n-by-p matrix, X, containing a Latin hypercube sample of n values
%on each of p variables. For each column of X, the n values are randomly
%distributed with one from each interval (0,1/n), (1/n,2/n), ..., (1-1/n,1), 
%and they are randomly permuted
%The resulting design matrix has values from 0 to 1.


X = lhsdesign(num_scens,p); 
% 
X(:,43) = 0; %We do not change the basal DNA synthesis rate, so we set all the changes for that column to be zero
filename = 'LHS_sensitivity_Sphase_constants_p21_Cdk2.csv';
writematrix(X,filename);


%We then recast the LHS sampling to a 2-fold change distributed through LHS
%on the log2 scale. First we go from 0 to 1 to -1 to 1, and then from 1/2
%to 2 as the final multiplicative factor.
param_mod_mat = 2.^((X - 0.5)*2); 

t_sphase_mat = zeros(num_scens,1);


parfor scenario_num = 1:num_scens
    
param = zeros(55,1);


%There are 51 parameters. Values from Heldt et al. are defined below


kSyE2f    = 0.03   ;    %AU/min  constitutive E2F synthesis
kSyE2fE2f = 0.04    ;   %AU/min  E2F-dependent E2F synthesis
jSyE2f    = 0.2      ;  %AU  Michealis-Menten constant for E2F synthesis
kAsRbE2f  = 5          ; %1/(AUmin)  association of Rb and E2F
kDsRbE2f  = 0.005     ;%1/min  dissociation of Rb:E2F complexes
kDeE2f    = 0.05    ; %1/min  constiutive E2F degradation

kPhRbCd   = 0.2     ;%1/(AUmin)  CycD:CDK4/6-mediated Rb phosphorylation
kPhRbCe   = 0.3      ; %1/(AUmin)  CycE:CDK2-mediated Rb phosphorylation
kPhRbCa   = 0.3      ;%1/(AUmin)  CycA:Cdk2-mediated Rb phosphorylation
kDpRb     = 0.05    ;   %1/min  dephosphorylation of Rb

kSyE1   = 0.005       ;  %1/min  constitutive Emi1 synthesis
kDeE1C1 = 0.005    ; %1/min  Cdh1-mediated Emi1 degradation
kDeE1   = 0.0005       ;%1/min  constitutive Emi1 degradation

kPhC1   = 0            ;%1/min  constitutive Cdh1 phosphorylation
kPhC1Ce = 0.01     ;%1/(AUmin)  CycE-mediated Cdh1 phosphorylation
kPhC1Ca = 1       ;%1/(AUmin)  CycA-mediated Cdh1 phosphorylation
kDpC1   = 0.05        ;%1/min  Cdh1 dephosphorylation
kAsE1C1 = 10         ;%1/(AUmin)  association of Emi1 and Cdh1
kDsE1C1 = 0.01       ;%1/min  dissociation of Emi1:Cdh1 complexes

kSyP21    = 0.002   ;   %AU/min  constitutive p21 synthesis
kSyP21P53 = 0.008     ;  %1/min  p53-dependent synthesis of p21
kDeP21    = 0.0025     ; %1/min  constitutive p21 degradation
kDeP21Cy  = 0.007       ;%1/(AUmin)  cyclin:CDK2-mediated p21 degradation (Skp2-dependent)
kDeP21aRc = 1           ;%1/(AUmin)  aRC-mediated p21 degradation (Cdt2-dependent)

kSyCe     = 0.01       ; %1/min  constitutive CycE synthesis
kSyCa     = 0.02        ;%1/min  constitutive CycA synthesis
kAsCyP21  = 1          ; %1/(AUmin)  association of cyclin:CDK2 and p21
kDsCyP21  = 0.05        ;%1/min  dissociation of cyclin:CDK2:p21 complexes
kDeCe     = 0.004       ;%1/min  constitutive CycE degradation
kDeCa     = 0.01        ;%1/min  constitutive CycA degradation
kDeCeCa   = 0.015       ;%1/(AUmin)  CycA:CDK2-mediated CycE degradation (Skp2-dependent)
kDeCaC1   = 2            ;%1/(AUmin)  Cdh1-mediated CycA degradation

kImPc     = 0.003       ; %AU/min  PCNA import into the nucleus
kExPc     = 0.006       ; %1/min   PCNA export from the nucleus

kPhRc     = 0.1          ;%1/min  cyclin:CDK2-mediated phosphorylation/priming of RCs
kDpRc     = 0.05         ;%1/min  dephosphorylation of pRCs
jCy       = 1.8          ;%AU  CDK2 threshold for the phosphorylation/priming of RCs
n       = 6            ;%-  hill coefficient for the phosphorylation/priming of RCs
kAsRcPc   = 0.01         ;%1/(AUmin)  association of phosphorylated/primed RCs and PCNA
kDsRcPc   = 0.001        ;%1/min  dissociation of pRC:PCNA complexes
kAsPcP21  = 100           ;%1/(AUmin)  association of PCNA and p21
kDsPcP21  = 0.01         ;%1/min  dissociation of PCNA:p21 complexes
kSyDna    = 0.0093       ;%1/min  DNA synthesis by aRCs

kSyP53    = 0.05         ;%AU/min  constitutive p53 synthesis
kDeP53    = 0.0005         ;%AU/min  DNA damage-dependent degradation of p53
jP53      = 0.01         ;%AU  inhibition constant of p53 degradation by DNA damage


kReDam    = 0.001        ;%1/min  p53-independent DNA damage repair
kReDamP53 = 0.005        ;%1/min  p53-dependent DNA damage repair
jDam      = 0.5         ;%AU  DNA damage threshold for repair


 Cd   = 0.65; 	%AU  relative CycD:Cdk4/6 level
tRb  = 5        ;     	%AU  total level of Rb
tC1  = 1         ;       %AU  total APC/C^Cdh1 level
Skp2 = 1          ;      %-  relative SCF^Skp2 level
Cdt2 = 1           ;     %-  relative CRL4^Cdt2 level


rDsRc     = 1; %Disassembly of DNA complexes at end of S phase


%There are 20 variables. Initial values from Heldt et al. defined below
y_init = zeros(1,20);

y_init(1,5) = 0.6; %tP21
y_init(1,6) = 0.5; %tCe

y_init(1,7) = 1.3; %tCa

y_init(1,12) = 0.5; %aPcna
y_init(1,14) = 1; %Rc


%Assign parameters to a vector and multiply it element-wise for each
%parameter change vector defined from sampling.
param(1)=kSyE2f;
param(2)=kSyE2fE2f;
param(3)=jSyE2f;
param(4)=kAsRbE2f;
param(5)=kDsRbE2f;
param(6)=kDeE2f;
param(7)=kPhRbCd;
param(8)=kPhRbCe;
param(9)=kPhRbCa;
param(10)=kDpRb;
param(11)=kSyE1;
param(12)=kDeE1C1 ;
param(13)=kDeE1 ;
param(14)=kPhC1 ;
param(15)=kPhC1Ce ;
param(16)=kPhC1Ca ;
param(17)=kDpC1;
param(18)=kAsE1C1;
param(19)=kDsE1C1;
param(20)=kSyP21 ;
param(21)=kSyP21P53 ;
param(22)=kDeP21 ;
param(23)=kDeP21Cy;
param(24)=kDeP21aRc;
param(25)=kSyCe ;
param(26)=kSyCa ;
param(27)=kAsCyP21 ;
param(28)=kDsCyP21;
param(29)=kDeCe ;
param(30)=kDeCa;
param(31)=kDeCeCa ;
param(32)=kDeCaC1;
param(33)=kImPc;
param(34)=kExPc ;
param(35)=kPhRc;
param(36)=kDpRc ;
param(37)=jCy;
param(38)=n ;
param(39)=kAsRcPc;
param(40)=kDsRcPc;
param(41)=kAsPcP21;
param(42)=kDsPcP21;
param(43)=kSyDna ;
param(44)=kSyP53 ;
param(45)=kDeP53 ;
param(46)=jP53;
param(47)=kReDam;
param(48)=kReDamP53 ;
param(49)=jDam ;

%Other global variables
param(50)=Cd;
param(51)=tRb;
param(52)=tC1;
param(53)=Skp2;
param(54)=Cdt2;
param(55)=rDsRc;

param = param.*param_mod_mat(scenario_num,:)';

%Define a tight grid of simulation time points - important to account for
%variable step sizes from the ode23s solver
tspan_1 = (0:0.1:5000);

%Solve
[t_1, y_1] = ode23s(@(t,y) detsys_Sphasetiming(t,y,param), tspan_1, y_init);

%Check whether simulation is proliferative or not
if y_1(end,18)>0.9
%If it is proliferative, calculate time at which S-phase entry happens
%To get accurate S-phase timing, we use linear interpolation to
%find the point at which the DNA amount is exactly 0.1.

    DNA_thres = 0.1;
    indnum = 18;
[~, t_sphase_ind] = min(abs(y_1(:,indnum) - DNA_thres));

t_sphase = interp1([y_1(t_sphase_ind-1,indnum) y_1(t_sphase_ind,indnum) y_1(t_sphase_ind+1,indnum)],[t_1(t_sphase_ind-1) t_1(t_sphase_ind) t_1(t_sphase_ind+1)],DNA_thres);


t_sphase_mat(scenario_num) = t_sphase;


end


end
%Save results for posterity
save('t_sphase_aCdk2_p21_sensitivity.mat','t_sphase_mat','X')

toc
 %% Sensitivity analysis with linear model fitting

%Parameter names
param_name{1}='kSyE2f';
param_name{2}='kSyE2fE2f';
param_name{3}='jSyE2f';
param_name{4}='kAsRbE2f';
param_name{5}='kDsRbE2f';
param_name{6}='kDeE2f';
param_name{7}='kPhRbCd';
param_name{8}='kPhRbCe';
param_name{9}='kPhRbCa';
param_name{10}='kDpRb';
param_name{11}='kSyE1';
param_name{12}='kDeE1C1';
param_name{13}='kDeE1';
param_name{14}='kPhC1';
param_name{15}='kPhC1Ce';
param_name{16}='kPhC1Ca';
param_name{17}='kDpC1';
param_name{18}='kAsE1C1';
param_name{19}='kDsE1C1';
param_name{20}='kSyP21';
param_name{21}='kSyP21P53';
param_name{22}='kDeP21';
param_name{23}='kDeP21Cy';
param_name{24}='kDeP21aRc';
param_name{25}='kSyCe';
param_name{26}='kSyCa';
param_name{27}='kAsCyP21';
param_name{28}='kDsCyP21';
param_name{29}='kDeCe';
param_name{30}='kDeCa';
param_name{31}='kDeCeCa';
param_name{32}='kDeCaC1';
param_name{33}='kImPc';
param_name{34}='kExPc';
param_name{35}='kPhRc';
param_name{36}='kDpRc';
param_name{37}='jCy';
param_name{38}='n';
param_name{39}='kAsRcPc';
param_name{40}='kDsRcPc';
param_name{41}='kAsPcP21';
param_name{42}='kDsPcP21';
param_name{43}='kSyDna';
param_name{44}='kSyP53';
param_name{45}='kDeP53';
param_name{46}='jP53';
param_name{47}='kReDam';
param_name{48}='kReDamP53';
param_name{49}='jDam';

%Other global variables
param_name{50}='Cd';
param_name{51}='tRb';
param_name{52}='tC1';
param_name{53}='Skp2';
param_name{54}='Cdt2';

param_name{55}='rDsRc';

%Fit a linear model with the Latin Hypercube terms as input and sphase
%entry time as output, but only for proliferative cells.
mdl = fitlm(X(t_sphase_mat~=0,:),t_sphase_mat(t_sphase_mat~=0));

param_name = reordercats(categorical(param_name),param_name);
 
%Calculate the slopes on the linear model for every parameter. This is the
%measure of sensitivity for the model
sens = mdl.Coefficients.Estimate(2:end);

bar(param_name, sens)

set(gca,'Fontsize',18)
set(gca,'LineWidth',2)
set(gca,'TickDir','out')
title('Sensitivity of S phase entry time to each parameter')
```
