## Supplementary material for "Inertial effect of cell state velocity on the quiescence-proliferation fate decision in breast cancer": Code_Inertia_p21Cdk2.zip: tall_landscapegen.m

```
%% This code will be used for generating the energy landscape of really large datasets

%This code makes use of the "tall" object type in MATLAB. This helps handle
%objects that do not fit in memory by sequential processing of the file
%split across multiple workers

%Initialize parallelization
parpool(8)

%Create a datastore that can access the csv file containing the
%trajectories
ds = datastore('gillespie_paths.csv');

%Set variable names
varnames = {'time' 'aPcna' 'pRc' 'aRc' 'Ca' 'P21' 'CaP21' 'Ce' 'CeP21' 'E1' 'C1' 'E1C1' 'iPcna' 'iRc' 'Rb' 'E2f' 'RbE2f' 'pC1' 'pRb' 'Rc' 'P53'};

ds.VariableNames = varnames;

%Convert to a tall data structure for handling large objects
traj = tall(ds);

% Set active Cdk2 and total p21 as A and B
A = traj.Ca + traj.Ce;
B = traj.P21 + traj.CaP21 + traj.CeP21 + traj.iRc + traj.iPcna;

%Probability calculation and landscape generation
h = histogram2(B, A, [50,50]);
h = h.Values;
h = h/sum(sum(h));
h_mod2 = h;
U = -log(h_mod2);
 U(~isfinite(U))=max(max(U(isfinite(U)))); %Sets infinite potentials to the highest existing potential

 %Interpolation for smoothing
xi = linspace(1, 50, 200);
yi = xi;
[xi, yi] = meshgrid(xi, yi);
x = (1:50);
y = (1:50);
Ui = interp2(x, y, U, xi, yi, 'spline');

xi = (gather(min(A)):gather(range(A))/199:gather(max(A)));
yi = (gather(min(B)):gather(range(B))/199:gather(max(B)));

%Saving the landscape variables if later analysis is required
save('Energylandscape.mat','xi','yi','Ui','U');

%Plot landscape
landscape = surf(xi, yi, Ui, 'LineStyle', 'none', 'FaceColor', 'interp', 'FaceAlpha', 0.9);
view([-45, 30])

%Save landscape figure
saveas(landscape,'output','fig');
```
